# Cyclin E/CDK2 deficiency links cell cycle control to intestinal barrier integrity and tight junction regulation

**DOI:** 10.64898/2026.10.12.758447

**Authors:** Hannes Hatten, Anna M. Verwaayen, Melissa Corcini-Berndt, Emiel P. C. van der Vorst, Daniela Paffen, Yulia A. Nevzorova, Sanne L. Maas, Christian Penners, Christian Liedtke

## Abstract

**Background & Aims:** The intestinal epithelium renews continuously through cell cycle-dependent turnover, yet it remains unknown whether core cell cycle regulators directly control intestinal barrier function. We investigated how loss of the Cyclin E1/Cyclin E2/CDK2 complex affects intestinal homeostasis. Based on the important role of this complex in compensatory liver regeneration, we hypothesized that its combined loss would impair epithelial renewal and barrier integrity.

**Methods:** We generated *Ccne2*^−/−^ mice and triple knockout (TKO) mice with intestinal epithelial cell (IEC)-specific deletion of *Ccne1* and *Cdk2* on a constitutive *Ccne2*-null background and characterized intestinal architecture, proliferation, and barrier function using histology, immunoblotting, qPCR, and FITC-dextran permeability assays. Mechanistic findings were validated in CRISPR/Cas9-generated *Ccne2*-deficient and CDK1-overexpressing MC38 colon carcinoma cells using transcriptomics and kinomics approaches.

**Results:** Contrary to our hypothesis, TKO maintained and *Ccne2*^−/−^ mice even enhanced cell cycle activity selectively in the colon along with enhanced *Cdk1* expression, had significantly lower serum FITC-dextran levels after oral administration, and increased *Tjp1*/ZO-1 expression, consistent with reduced intestinal permeability. *Ccne2*-deficient MC38 cells recapitulated the increase in ZO-1 and exhibited transcriptional enrichment of gene sets associated with tight junctions and cell cycle. CDK1 overexpression in MC38 cells increased ZO-1 protein abundance and was associated with activation of an AMPK-related kinase network. In TKO and *Ccne2*^−/−^ colonic epithelium, CDK1, phosphorylated AMPK and ZO-1 showed partial colocalization at lateral epithelial cell-cell contacts.

**Conclusions:** Loss of Cyclin E2, either alone or in combination with epithelial Cyclin E1/CDK2 deficiency, is associated with increased CDK1 expression and strengthens the colonic barrier integrity in mice. Findings from MC38 cells suggest that transcriptional reprogramming and enhanced CDK1 signaling in the context of Cyclin E2 deficiency may contribute to tight junction reinforcement and improved barrier function.

## Introduction

The intestinal epithelium is one of the most rapidly self-renewing tissues in the mammalian body, undergoing complete turnover every 3-5 days [1]. This continuous renewal is driven by actively cycling intestinal stem cells (ISCs) residing at the base of the crypts, which can be identified by expression of the WNT target gene *Lgr5* [2]. These crypt base columnar cells are interspersed among Paneth cells, which provide essential niche signals, including WNT3, EGF, and Notch ligands, and divide approximately once every 24 hours generating a pool of highly proliferative transit-amplifying progenitors [3]. As TA cells migrate upward along the crypt-villus axis, they progressively exit the cell cycle, terminally differentiate into the specialized epithelial lineages (enterocytes, goblet cells, enteroendocrine cells, tuft cells), and are ultimately shed into the lumen at the villus tip [1]. This tightly coordinated balance between proliferation, differentiation, and cell loss is a prerequisite for epithelial homeostasis, and its dysregulation is a hallmark of intestinal pathologies ranging from inflammatory bowel disease to colorectal carcinogenesis.

At the molecular level, cell cycle progression in the crypt compartment is governed by sequential activation of cyclin-dependent kinases (CDKs) in complex with their regulatory cyclin subunits. Canonical Wnt/β-catenin signaling, which is constitutively active in the crypt base, drives expression of key cell cycle activators such as cyclin D1 and c-Myc, thereby promoting ISCs and TA cells to enter and progress through G1 phase [4]. In turn, Notch signaling maintains the proliferative crypt compartment in part by repressing the CDK inhibitors p27Kip1 and p57Kip2 [5]. Despite this detailed understanding of the upstream signals that promote proliferation, comparatively little is known about the specific contribution of the core G1/S cell cycle machinery to intestinal epithelial renewal. However, pharmacological inhibition of CDK activity in intestinal epithelial cells (IECs) has been shown to trigger DNA damage responses, cell cycle arrest, and apoptosis, suggesting that CDK activity is required to maintain epithelial turnover [6]. Given that disturbed epithelial proliferation and differentiation can compromise gut barrier integrity and secondarily affect hepatic homeostasis *via* the gut-liver axis, a more detailed understanding of the core cell cycle machinery operating in IECs is of considerable pathophysiological interest.

Cyclin E1 and Cyclin E2, together with their catalytic partner CDK2, orchestrate the G1/S transition, yet their specific contributions to intestinal epithelial biology remain incompletely understood. Cyclin E/CDK2 complexes phosphorylate substrates including the retinoblastoma protein (Rb), and components of the pre-replication complex, thereby releasing E2F-dependent transcription and facilitating the onset of DNA replication [7]. In many tissues, Cyclin E1 and Cyclin E2 are considered largely functionally redundant, and constitutive inactivation of each individual gene is compatible with grossly normal development and tissue homeostasis, suggesting that compensatory mechanisms may mask the individual contribution of each cyclin to G1/S control. Our previous work in the liver showed that single genetic deletion of either *Ccne1*, *Ccne2* or *Cdk2* in mice has, at best, moderate effects on liver homeostasis and regeneration. However, concomitant loss of all three components resulted in severely impaired hepatocyte proliferation and liver mass restoration following partial hepatectomy [8, 9]. Besides that, we previously demonstrated that loss or inhibition of *Ccne1* is largely protective, as it prevents the onset and progression of experimental liver fibrosis [10], hepatic acute inflammation [11, 12], and hepatocellular carcinoma [13].

However, the functional relevance of E-type cyclins and CDK2 in the gut epithelium remains unclear. Moreover, alterations in epithelial proliferation and differentiation can profoundly affect intestinal barrier integrity, which is governed largely by tight junction (TJ) complexes and is essential for preventing translocation of luminal antigens and microbial products into the lamina propria and, ultimately, the portal circulation [14]. This barrier function is of particular relevance in the context of the gut-liver axis, since the liver is continuously exposed *via* the portal vein to gut-derived antigens and metabolites, and increased intestinal permeability has been linked mechanistically to a range of hepatic pathologies, including fatty liver disease, fibrosis, and cirrhosis [15]. Interestingly, recent work by Kuo *et al.* has challenged the conventional view that individual TJ proteins act merely as static, structural components of the barrier. Using mice with intestinal epithelial-specific deletion of *Tjp1*, which encodes ZO-1, the authors demonstrated that ZO-1 is largely dispensable for baseline TJ barrier function but is instead specifically required for the proliferative response of the epithelium during mucosal repair after injury [16]. These findings reveal a previously unrecognized, non-canonical link between TJ components and epithelial proliferation and conversely raise the possibility that regulators of epithelial cell cycle progression may, in turn, influence TJ composition and barrier integrity.

Given the central role of the Cyclin E/CDK2 complexes in cell cycle control, we hypothesized that loss of E-type cyclins and CDK2 in IECs would alter epithelial turnover and, thereby, influence barrier function, with potential downstream effects for the gut-liver axis. Here, we generated triple knockout (TKO) mice with IEC-specific deletion of *Ccne1* and *Cdk2* on a constitutive *Ccne2*-null background and performed a comprehensive analysis of epithelial morphology, proliferation, barrier integrity, and downstream effects on liver homeostasis. Our findings reveal an unexpected association between Cyclin E/CDK2 deficiency, intestinal barrier regulation, and modulation of TJ components.

## Results

### **Expression pattern of E**-**type cyclins and** *Cdk2* **along the murine intestine**

As an initial step, we assessed the expression of *Ccne1*, *Ccne2*, and *Cdk2* across the three anatomical segments of the murine small intestine (proximal: duodenum; medial: jejunum; distal: ileum), in small-intestinal (SI) organoids derived from these regions, and in colon tissue. Expression values were normalized to primary resting wildtype (WT) hepatocytes, which express *Cdk2* constitutively but lack relevant levels of E-type cyclins [8]. SI-derived organoids showed significantly higher expression of *Ccne1* and *Ccne2*, whereas *Cdk2* levels were comparable to resting hepatocytes, suggesting that E-type cyclins contribute to organoid proliferation (Figure 1A-C). In native intestinal tissue, *Ccne1* expression was only moderately detectable, with a slight but significant increase in distal SI and colon, compared with resting hepatocytes (Figure 1A). In contrast, *Ccne2* expression was substantially elevated across all intestinal segments (Figure 1B), while *Cdk2* levels were lower or similar compared with resting hepatocytes (Figure 1C). Together, these data indicate a segment-specific regulation of E-type cyclins along the gut and suggest that Cyclin E2 may play a particularly prominent role in intestinal epithelial homeostasis.

**Figure 1.**
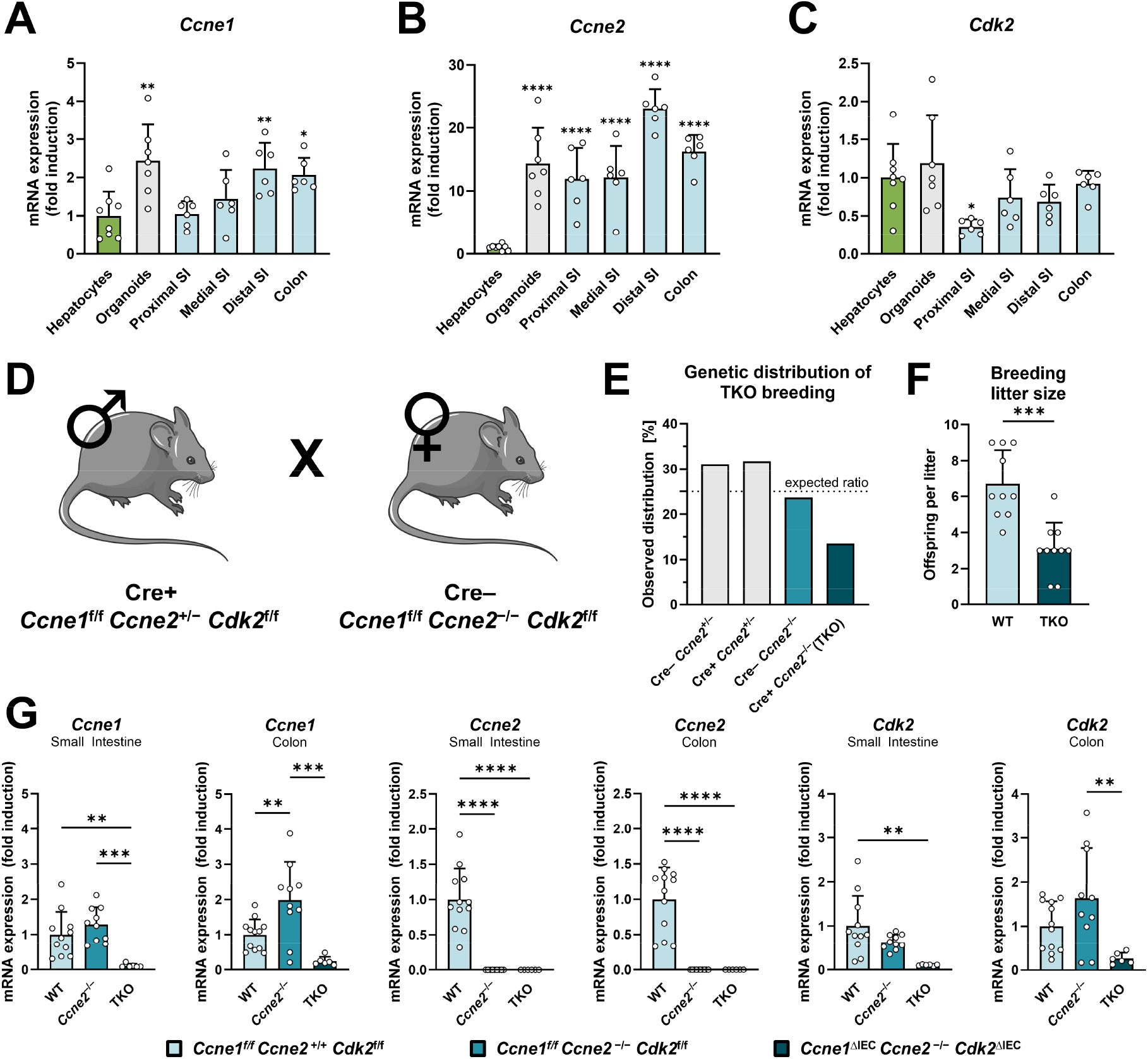
Expression pattern of E-type cyclins and *Cdk2* along the murine intestine. The mRNA expression profiles of (**A**) *Ccne1*, (**B**) *Ccne2* and (**C**) *Cdk2* in intestinal organoids derived from small intestine (SI), as well as in the proximal, medial and distal parts of SI and colon of 12-week-old WT mice were investigated by quantitative real-time PCR (qPCR). Data was normalized against the expression of the housekeeping gene *Gapdh* and calculated as fold induction compared to isolated resting primary WT hepatocytes. All bars indicate the outcome of at least n = 6 individual samples per group, statistical significance was calculated in comparison to hepatocytes, significance is indicated above respective bars. (**D**-**F**) Triple-knockout (TKO) mice with IEC-specific deletion of Cyclin E1 and CDK2 and constitutive deletion of Cyclin E2 (*Ccne1*^ΔIEC^ *Ccne2*^−/−^ *Cdk2*^ΔIEC^) were generated and analyzed at the age of 12 weeks. TKO mice were compared to constitutive Cyclin E2 knockout mice (*Ccne1*^f/f^ *Ccne2*^−/−^ *Cdk2*^f/f^) and WT mice (*Ccne1*^f/f^ *Ccne2*^+/+^ *Cdk2*^f/f^). (**D**) Exemplary mating scheme: Male mice expressing Cre recombinase under control of the Villin 1 promotor (Cre+) and heterozygous for Cyclin E2 (*Ccne1*^f/f^ *Ccne2*^+/−^ *Cdk2*^f/f^) were crossed with Cre-negative, female mice constitutively lacking Cyclin E2 (*Ccne1*^f/f^ *Ccne2*^−/−^ *Cdk2*^f/f^). (**E**) Observed genetic distribution of offspring from TKO breeding couples. Please note that in case of Mendelian distribution 25% of the offspring (dashed line) should have a TKO genotype. Genotypes of 150 offspring mice from TKO breeding couples were determined by polymerase chain reaction (PCR). (**F**) Litter size of TKO breeding couples compared to WT breeding couples is shown. (**G**) Gene expression of *Ccne1*, *Ccne2* and *Cdk2* in small intestine and colon of WT, *Ccne2*^−/−^ and TKO mice was investigated by quantitative real-time PCR (qPCR). Data were normalized against the expression of the housekeeping gene *Gapdh* and calculated as fold induction compared to WT controls. All bars represent the outcome of at least n = 6 individual samples per group. Data are presented as mean ± SD; one-way ANOVA followed by Dunnett’s multiple comparisons test (**A**-**C**) or Tukey’s multiple comparisons test (**G**) and unpaired two-tailed *t* test (**F**) were used for statistical analysis; Significances were defined as **: *p* < 0.01; ***: *p* < 0.001; ****: *p* < 0.0001.

### Generation and breeding characteristics of intestinal-epithelial cell-specific Cyclin E1/E2/Cdk2 triple-knockout mice

We generated mice with IEC-specific deletion of *Ccne1* and *Cdk2* on a constitutive *Ccne2*-null background (*Ccne1*^ΔIEC^ *Ccne2*^−/−^ *Cdk2*^ΔIEC^; TKO) using Villin1-Cre-driven recombination [17] and previously described floxed alleles of *Ccne1* and *Cdk2* [8, 18–20]. As *Ccne2*^−/−^ male mice are usually sterile [18], we applied the breeding strategy shown in Figure 1D. Genotyping of offspring from TKO breeding pairs revealed a lower-than-expected proportion of TKO animals relative to the Mendelian ratio (Figure 1E). In addition, TKO breeding pairs produced significantly smaller litters than WT controls (Figure 1F), suggesting a partial reduction in perinatal viability and/or fertility associated with the TKO genotype.

Despite this early disadvantage, TKO mice exhibited normal postnatal growth and no overt macroscopic abnormalities. qPCR analysis revealed an overall reduction in *Ccne1*, *Ccne2*, and *Cdk2* transcript levels in the small intestine and colon of TKO mice compared with WT controls (Figure 1G). Of note, some comparisons did not reach statistical significance, which may reflect residual expression of *Ccne1* and *Cdk2* in non-epithelial (and therefore Cre-negative) cells. Notably, *Ccne1* expression was significantly upregulated in the colon of *Ccne2*^−/−^ mice, consistent with compensatory regulation previously observed in our earlier studies of the liver [9].

### Loss of Cyclin E/CDK2 in IECs elongates villus-crypt units and selectively increases cell-cycle activity in the colon

To determine whether IEC-specific loss of Cyclin E/CDK2 affects intestinal architecture, we compared morphological parameters in WT, *Ccne2*^−/−^ and TKO mice. Body weight and intestinal length were unchanged across all genotypes (Figure 2A).

**Figure 2:**
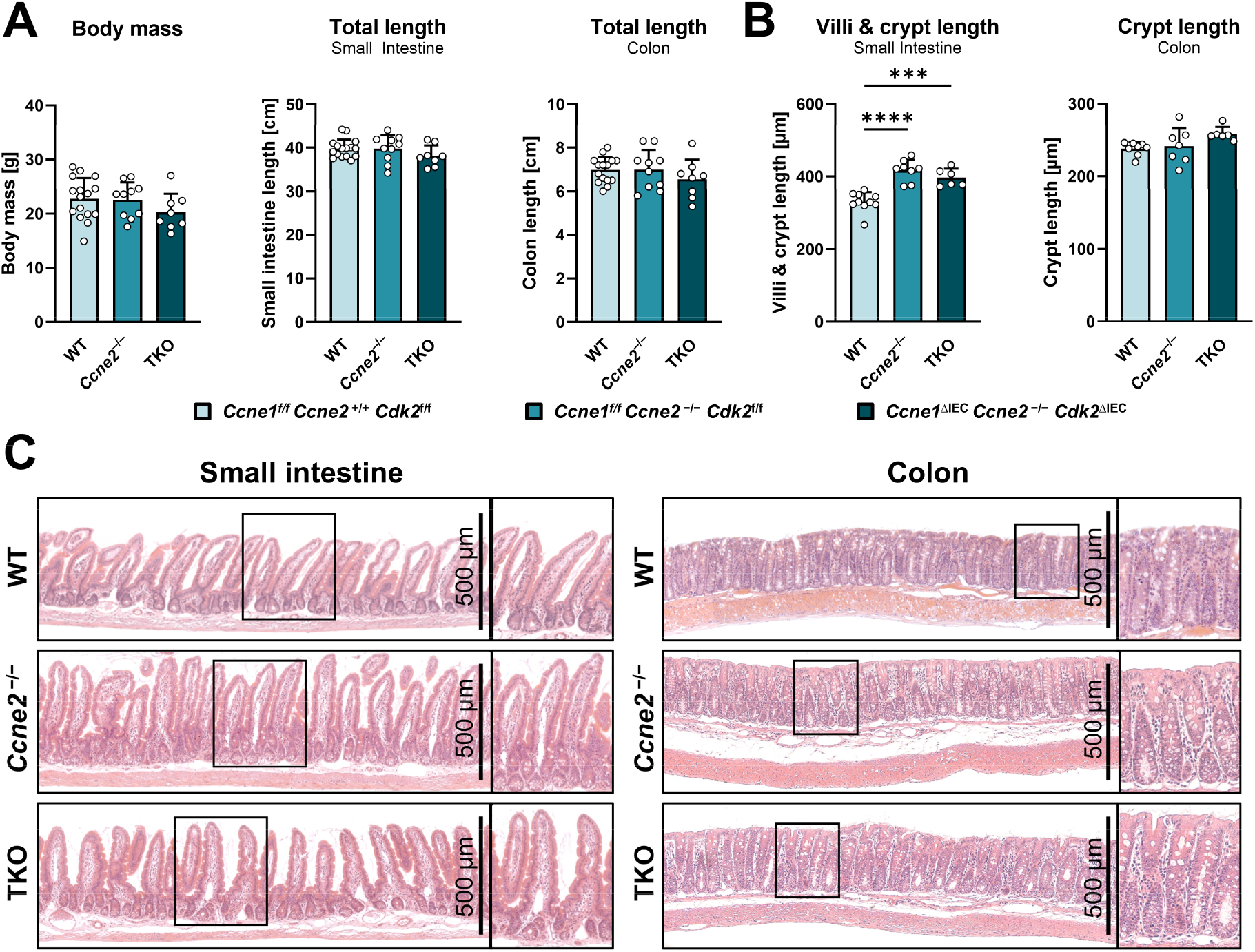
Deletion of Cyclin E and CDK2 in IECs does not result in a macroscopic phenotype but leads to villi and crypt elongation. Small intestine and colon of *Ccne2*^−/−^ and TKO mice were analyzed at the age of 12 weeks regarding morphological and histological alterations in comparison to WT mice. (**A**) Analysis of body mass, length of the small intestine and colon. (**B**) The length of crypt and villi was determined by measuring the distance from crypt base to villus tip. The distance was measured at ten different points along the tissue section for each mouse and plotted as the mean value. (**C**) Representative Hematoxylin and Eosin (H&E) stained small intestine (left) and colon (right) sections of WT, *Ccne2*^−/−^ and TKO mice. Boxed areas are depicted at the right part of the image as enlarged view with 1.2-fold (left) and 2.0-fold (right) magnifications. All bars indicate the outcome of at least n = 6 individual samples per group. Data are presented as mean ± SD; one-way ANOVA followed by Tukey’s multiple comparisons test was used for statistical analysis (**A**, **B**); Significances were defined as ***: *p* < 0.001; ****: *p* < 0.0001.

A more detailed histological analysis revealed that villus-crypt units in the SI were on average approximately 80µm longer in both TKO and *Ccne2*^−/−^ mice compared with WT controls (Figure 2B-C). In the colon, crypt length was at best mildly increased in TKO mice compared to the controls. Aside from these changes, Hematoxylin and Eosin (H&E) staining did not reveal additional abnormalities (Figure 2C).

The Cyclin E/CDK2 complex governs the G1/S transition and thus represents a central node of the cell cycle machinery. We therefore assessed how IEC-specific loss of Cyclin E/CDK2 affects cell cycle activity *in vivo*. Immunohistochemical staining for the proliferation marker Ki-67 was performed on small intestinal and colonic tissue sections. As expected for intestinal crypt architecture, Ki-67-positive cells were concentrated at the crypt base, with numbers declining progressively toward the crypt-villus axis as cells differentiate (Figure 3A). Only few Ki-67-positive cells were detected outside the epithelium, in the lamina propria or submucosa. No overt differences in the distribution or abundance of Ki-67^+^ cells were observed among WT, *Ccne2*^−/−^ and TKO mice.

**Figure 3:**
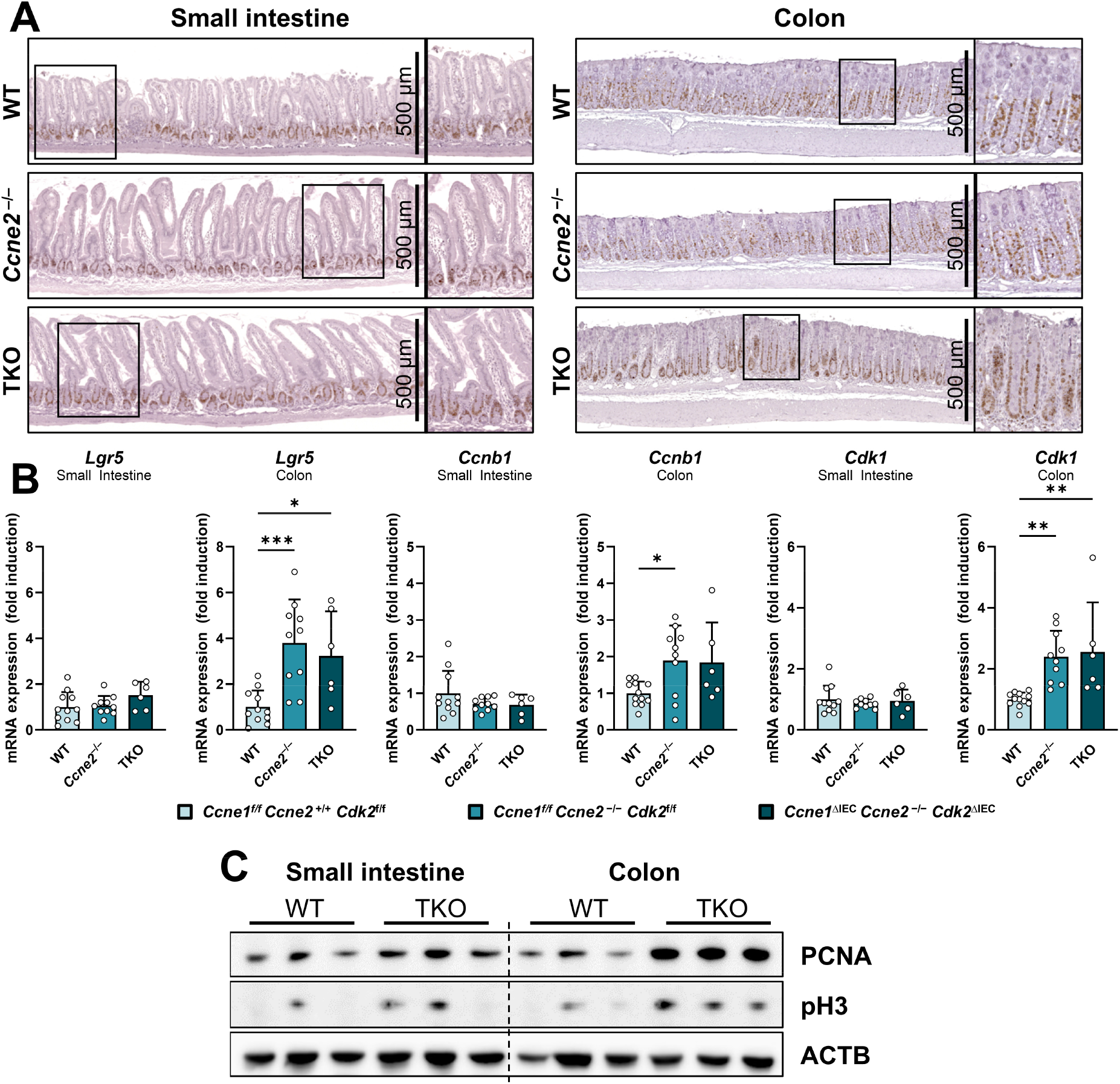
Deletion of Cyclin E and CDK2 in IECs increases overall cell cycle activity in the colon. Small intestine and colon of *Ccne2*^−/−^ and TKO mice were analyzed at the age of 12 weeks regarding cell cycle activity in comparison to WT mice. (**A**) Representative Ki-67 nuclear stainings (brown) of small intestine (left) and colon (right) sections. Boxed areas are depicted at the right part of the image as enlarged view with 1.2-fold (left) and 2.0-fold (right) magnifications. (**B**) Gene expression of *Lgr5*, *Ccnb1* and *Cdk1* in small intestine and colon was investigated by quantitative real-time PCR (qPCR). Data were normalized against the expression of the housekeeping gene *Gapdh* and calculated as fold induction compared to WT controls. All bars indicate the outcome of at least n = 6 individual samples per group. (**C**) Immunoblot analysis of PCNA and pH3 in small intestine and colon protein extracts from TKO and WT mice (n = 3). Abundance of ACTB was determined as an internal loading control. Data are presented as mean ± SD; one-way ANOVA followed by Tukey’s multiple comparisons test was used for statistical analysis (**B**); Significances were defined as *: *p* < 0.05; **: *p* < 0.01; ***: *p* < 0.001.

Because reliable quantification of Ki-67^+^ cells in intestinal tissue is inherently difficult, we assessed cell cycle activity using complementary molecular markers. qPCR analysis revealed that significant expression changes in selected cell cycle-associated genes were largely restricted to the colon, whereas no comparable alterations were detected in the small intestine (Figure 3B). Specifically, *Lgr5* and *Cdk1* expression was significantly increased in the colon of both *Ccne2*^−/−^ and TKO mice, whereas *Ccnb1* expression was significantly increased only in the colon of *Ccne2*^−/−^ mice. Together, these findings point to a colon-specific increase in cell cycle-associated gene expression following disruption of Cyclin E function. To further characterize cell cycle activity, we performed immunoblot analysis in WT and TKO tissue (Figure 3C). PCNA (S-phase marker) and phospho-histone H3 (pH3; M-phase marker) levels were comparable between genotypes in the small intestine, whereas both markers were markedly increased in the TKO colon relative to WT, indicating enhanced S- and M-phase activity specifically in this tissue. Together, these findings suggest a selective increase in cell cycle activity within the colon and provide complementary molecular evidence for enhanced proliferative signaling following combined loss of Cyclin E/CDK2 function.

### Loss of Cyclin E/CDK2 reduces intestinal permeability and basal hepatic *Tlr4* **expression**

Alterations of the intestinal regeneration and homeostasis can affect the intestinal barrier function [21]. To assess whether Cyclin E/CDK2 loss affects intestinal barrier function, we performed FITC-dextran assays in WT, *Ccne2*^−/−^ and TKO mice (Figure 4A). Both *Ccne2*^−/−^ and TKO mice exhibited significantly reduced serum FITC-dextran levels compared with WT controls (Figure 4B), consistent with reduced intestinal permeability. qPCR analysis revealed increased *Tjp1* expression in colon of *Ccne2*^−/−^ mice as well as in the colon and SI of TKO mice (Figure 4C). Immunofluorescence staining confirmed enhanced ZO-1 localization at apical junctions in colonic epithelium of *Ccne2*^−/−^ and TKO mice (Figure 4D).

**Figure 4:**
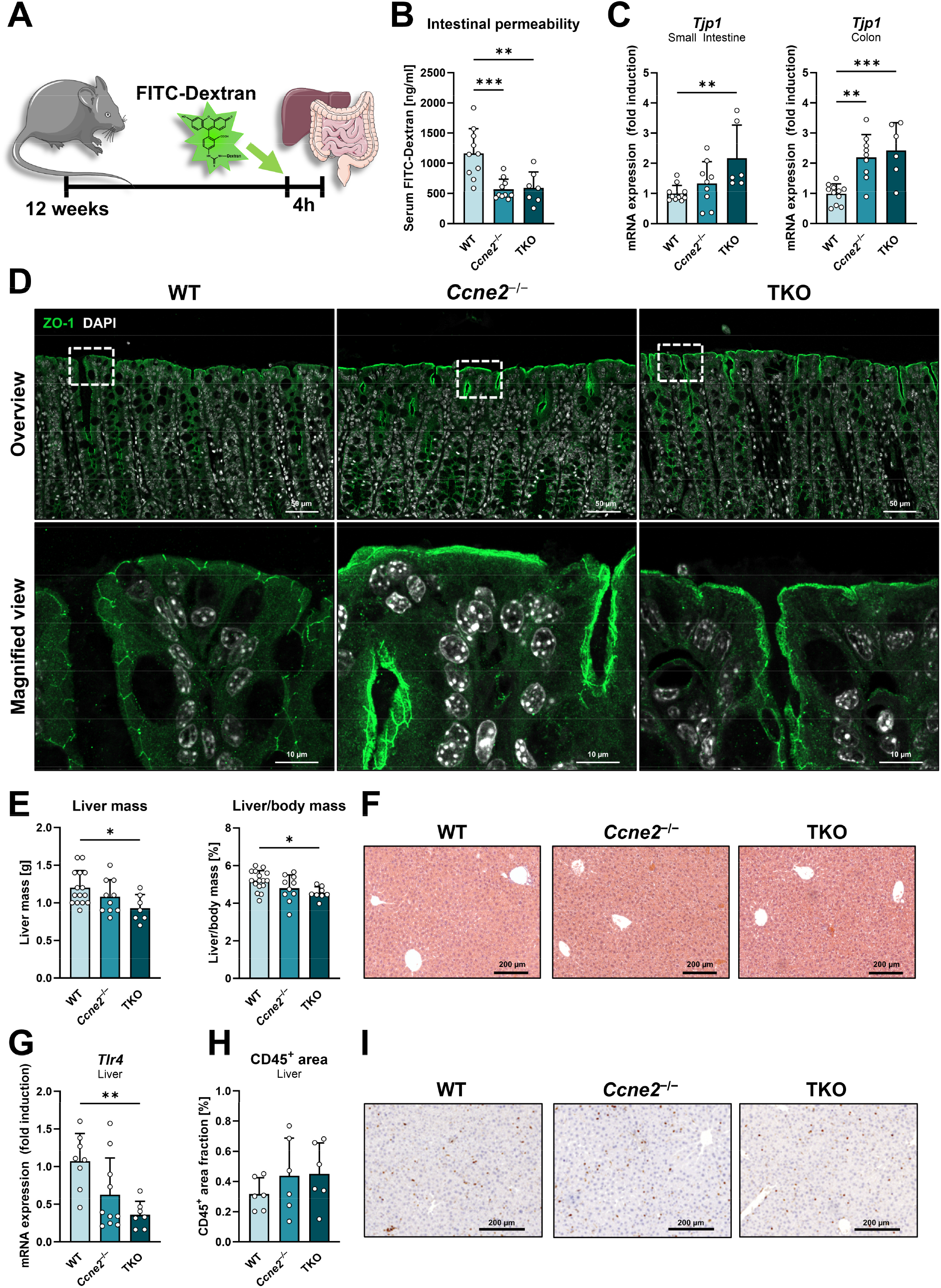
Deletion of Cyclin E/CDK2 in IECs enhances the intestinal barrier and reduces basal *Tlr4* gene expression in the liver. (**A**) Experimental setting. The gut barrier of 12-week-old *Ccne2*^−/−^ and TKO mice was investigated and compared to WT mice. Four hours before sacrifice, all experimental animals were given an oral dose of FITC-dextran to determine intestinal permeability. (**B**) Serum concentrations of FITC-conjugated dextran were determined spectrophotometrically 4h after oral administration to determine intestinal permeability. (**C**) Gene expression of *Tjp1* in small intestine and colon was investigated by quantitative real-time PCR (qPCR). (**D**) Representative confocal immunofluorescence stainings for ZO-1 (green) on paraffin sections of colon tissue from WT, *Ccne2*^−/−^ and TKO mice. Nuclei are stained with DAPI (white). Lower panels show magnified views of regions indicated by dashed boxes in the corresponding upper panels. (**E-I**) Livers of 12-week-old *Ccne2*^−/−^ and TKO mice were analyzed for morphological alterations and markers of microbial translocation in comparison to WT mice. (**E**) Left: Analysis of liver mass. Right: Liver mass index was calculated as the liver mass/ body mass ratio in percent. (**F**) Representative Hematoxylin and Eosin (H&E) stained liver sections of WT, *Ccne2*^−/−^ and TKO mice. (**G**) Gene expression of *Tlr4* in the liver was investigated by quantitative real-time PCR (qPCR). (**H**) Paraffin liver sections from WT, *Ccne2*^−/−^ and TKO mice were stained with a CD45-specific antibody. CD45^+^ image areas were quantified using ImageJ and calculated as percent of total tissue area. (**I**) Representative immunohistochemical stainings of CD45 (brown) on liver sections. Gene expression data were normalized against the expression of the housekeeping gene *Gapdh* and calculated as fold induction compared to WT controls. All bars indicate the outcome of at least n = 6 individual samples per group. Data are presented as mean ± SD; one-way ANOVA followed by Tukey’s multiple comparisons test was used for statistical analysis (**B**, **C**, **E**, **G**, **H**); Significances were defined as *: *p* < 0.05; **: *p* < 0.01; ***: *p* < 0.001.

Given the role of the intestinal barrier in limiting microbial translocation, we next examined the respective liver tissues for changes in morphological, histological and anti-inflammatory parameters. TKO mice displayed significantly reduced liver mass and liver-to-body-mass ratio (Figure 4E), without evidence of pathological changes on histological examination (Figure 4F). qPCR analysis showed reduced hepatic *Tlr4* expression in TKO mice and a trend towards reduced expression (*p* = 0.059) in *Ccne2*^−/−^ mice (Figure 4G). This finding is consistent with altered gut-liver signaling, potentially involving LPS-TLR4 pathways. Of note, the number of hepatic CD45-positive immune cells did not differ between liver sections of WT, *Ccne2*^−/−^ and TKO mice (Figure 4H-I) indicating that the reduced hepatic *Tlr4* expression is not associated with a diminished number of immune cells *per se*.

Taken together, *Ccne2*^−/−^ and TKO mice exhibit reduced intestinal permeability, accompanied by increased colonic *Tjp1* expression and junctional ZO-1 localization, alongside increased colonic expression of *Cdk1* and *Ccnb1*. Reduced hepatic expression of *Tlr4* in TKO mice indicates that Cyclin E/CDK2 deletion may reflect altered gut-liver signaling and potentially reduced microbial translocation into the liver under basal conditions.

### Loss of Cyclin E2 in MC38 cells increases ZO-1 expression and alters cell cycle progression without affecting cell expansion

From the previous *in vivo* results, we hypothesized that loss of Cyclin E2 in colon may result in a compensatory over-activation of the intestinal cell cycle machinery involving *Cdk1*, and *Ccnb1* which in turn triggers up-regulation of *Tjp1*/ZO-1 and potentially other TJ related factors through yet unknown mediators and pathways (Figure 5A).

**Figure 5:**
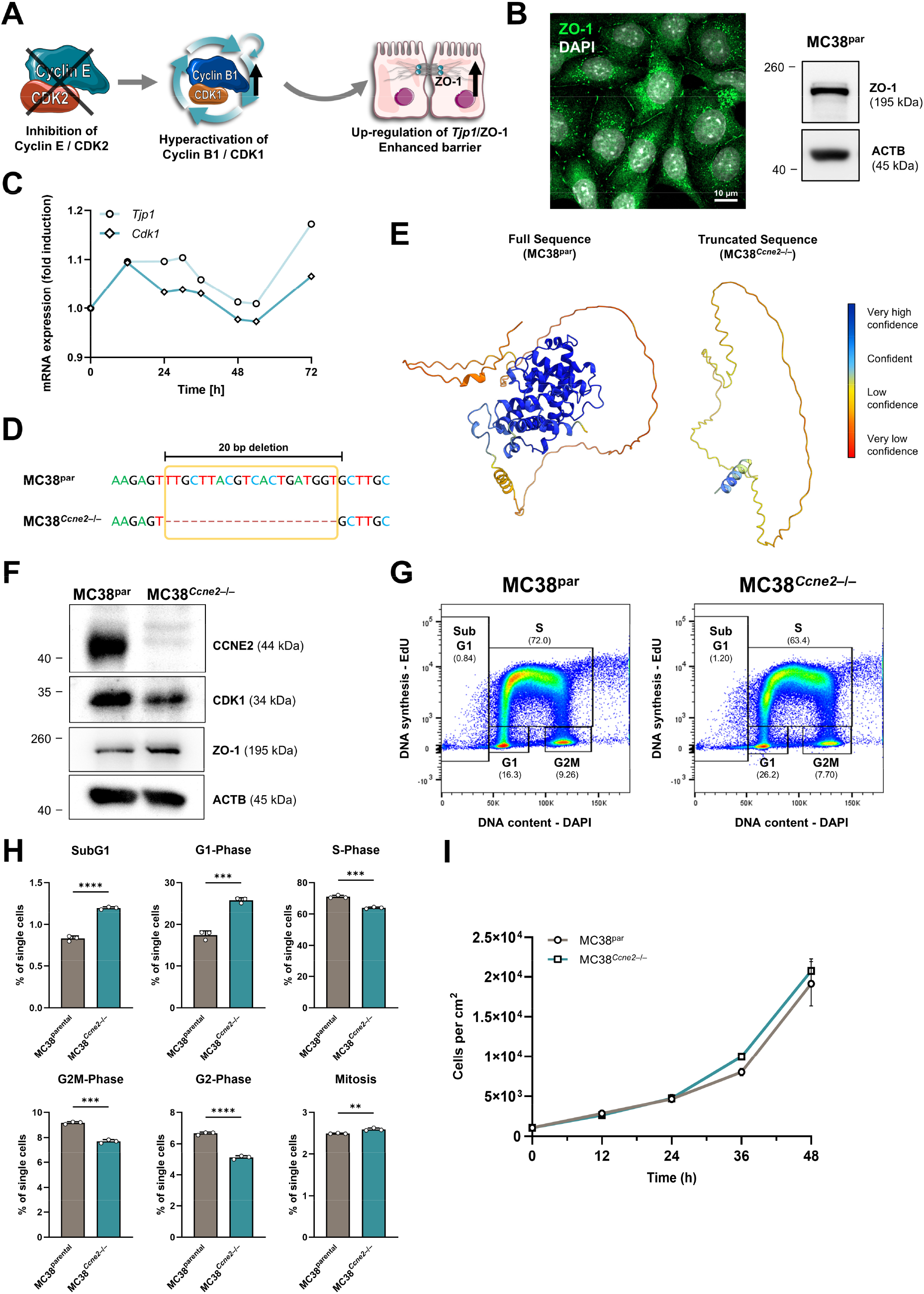
Loss of Cyclin E2 in MC38 cells increases ZO-1 expression and alters cell cycle progression without affecting cell expansion. (**A**) Schematic representation of the proposed working hypothesis. Loss of Cyclin E2 is hypothesized to induce compensatory activation of the Cyclin B1-CDK1 axis, resulting in increased *Tjp1*/ZO-1 expression and potentially broader regulation of TJ-associated factors (**B**) Expression of ZO-1 in MC38 cells was assessed by confocal immunofluorescence microscopy (left; ZO-1: green, DAPI: white) and Western blot analysis (right) to evaluate suitability of MC38 cells as a model for investigation of the proposed mechanism. (**C**) Continuously proliferating MC38 cells were collected regularly over 72 hours and relative mRNA expression of *Cdk1* and *Tjp1* was determined by qPCR (n = 1). Expression was normalized to *Gapdh* and is shown as fold induction compared to timepoint 0 h. (**D**) Sequence alignment of the sgRNA-targeted region within the *Ccne2* coding sequence in parental MC38 cells (MC38^par^) and CRISPR/Cas9 engineered mutant cells (MC38^Ccne2–/−^) confirmed a 20-bp deletion, resulting in a frameshift and a predicted premature termination codon. (**E**) *In silico* structural predictions of wildtype murine Cyclin E2 and the mutant translation product, using AlphaFold2, illustrating the structural disruption expected in the *Ccne2* mutant. For direct structural comparison, the predicted truncated protein was displayed in the corresponding orientation of the wildtype protein. (**F**) Western blot of proteins from MC38^par^ and MC38*^Ccne2–/−^* cells probed for ZO-1, Cyclin E2 and CDK1. ACTB was probed as internal loading control. **(G-H**) MC38^par^ and MC38*^Ccne2–/−^* cells were collected after culture for 24 h followed by analysis using flow cytometry. (**G**) Representative cell cycle gating strategy of parental MC38 cells and MC38*^Ccne2^*^−/−^ cells based on EdU incorporation and cellular DNA content as measured by DAPI staining to discriminate between cell cycle phases. (**H**) Quantification of cell cycle phases (n = 3). (**I**) Cell growth kinetics analysis of MC38^par^ and MC38*^Ccne2–/−^* cells over a 48 h period (n = 3). Data are presented as mean ± SD; unpaired two-tailed *t* test (**H**) and 2way ANOVA followed by Šídák’s multiple comparisons test (**I**) was used for statistical analysis; Significances were defined as **: *p* < 0.01; ***: *p* < 0.001; ****: *p* < 0.0001.

To test the hypothesis that *Tjp1* is a cell cycle-regulated gene, we took advantage of murine MC38 colon carcinoma cells. We confirmed that MC38 cells express *Tjp1*/ZO-1 (Figure 5B). In continuously growing cells, we determined the gene expression profile of key cyclins (*i.e. Ccne1*, *Ccne2*, *Ccna2*, *Ccnb1*), CDKs (*i.e. Cdk1*, *Cdk2*) and *Tjp1* for a period of 72 hours. Interestingly, we found a highly significant correlation between the expression of *Tjp1* and *Cdk1* (*r* = 0.928, Figure 5C, Supplementary Figure 1A), a moderate correlation between the expression of both E-cyclins and *Tjp1*, but no correlation between *Tjp1* and *Ccna2* or *Ccnb1,* respectively (Supplementary Figure 1A-B).

To mimic the *Ccne2* gene deletion used in our *in vivo* studies, we applied CRISPR/Cas9 technology and generated a 20 bp frameshift deletion (c.685_704del with regard to mRNA-sequence NM_001282943.2) within the 5’ coding region of the *Ccne2* gene in MC38 cells Figure 5D). The c.685_704del mutation causes a frameshift beginning at codon 210 and introduces a premature termination codon after three aberrant amino acids. The predicted mutant protein comprises 212 amino acids instead of the 404 amino acids of wildtype Cyclin E2, resulting in aberrant three-dimensional structure, loss of most of the cyclin-box region and the complete C-terminal cyclin domain as predicted by Alphafold analysis [22](Figure 5E). The mutation is therefore predicted to represent a loss-of-function allele. Accordingly, Cyclin E2 protein was no longer detectable by immunoblot analysis in *Ccne2*-deficient MC38 cells (MC38*^Ccne2–/−^*) in comparison to parental (MC38^par^) cells (Figure 5F). Of note, MC38*^Ccne2–/−^* cells expressed slightly less CDK1, but more ZO-1 protein compared to MC38^par^ cells (Figure 5F), which in part phenocopies our observations in colon of *Ccne2*^−/−^ mice.

Comparative EdU incorporation and DNA-content analysis of MC38^par^ and MC38*^Ccne2–/−^* cells by flow cytometry revealed an increased proportion of sub-G1 cells in MC38*^Ccne2–/−^* cells, suggesting enhanced apoptotic susceptibility (Figure 5G-H). In parallel, MC38*^Ccne2–/−^* cells displayed reduced frequencies of S-phase and G2/M-phase cells, indicating impaired cell cycle progression through DNA replication and mitosis. Notably, these changes were accompanied by a relative accumulation of cells in G1, consistent with a partial delay at the G1/S transition. Despite these alterations in cell cycle dynamics, growth kinetic analysis over 48 h demonstrated comparable expansion of MC38^par^ and MC38*^Ccne2–/−^* cells (Figure 5I), indicating that Cyclin E2 is dispensable for short-term proliferation under standard culture conditions. Collectively, these findings suggest that Cyclin E2 facilitates efficient cell cycle progression but that compensatory mechanisms maintain overall proliferative capacity in its absence.

### Transcriptomic profiling reveals up-regulation of TJ and cell cycle-related genes due to loss of *Ccne2* in MC38 cells

To identify molecular pathways altered by Cyclin E2 loss, we performed 3′ mRNA-sequencing (mRNA-seq) in parental and *Ccne2*-deficient MC38 cells. Principal component analysis (PCA) of both transcriptomes revealed distinct clustering of MC38^par^ and MC38*^Ccne2–/−^* cells (Figure 6A). Based on an adjusted *p*-value < 0.05, 1,278 genes were upregulated, and 1,378 genes were downregulated in MC38*^Ccne2–/−^* cells compared to MC38^par^ (Figure 6B). Over-representation analysis of significantly upregulated genes on KEGG pathways revealed that genes related to “Tight junction” and “Cell cycle”, followed by “Apoptosis”, “Adherens junction”, “DNA replication” and “Colorectal cancer” were highly over-represented (Figure 6C); “Base excision repair”, “Mismatch repair” and “p53 signaling pathway” and some more pathways were also significantly enriched, albeit with lower gene counts.

**Figure 6:**
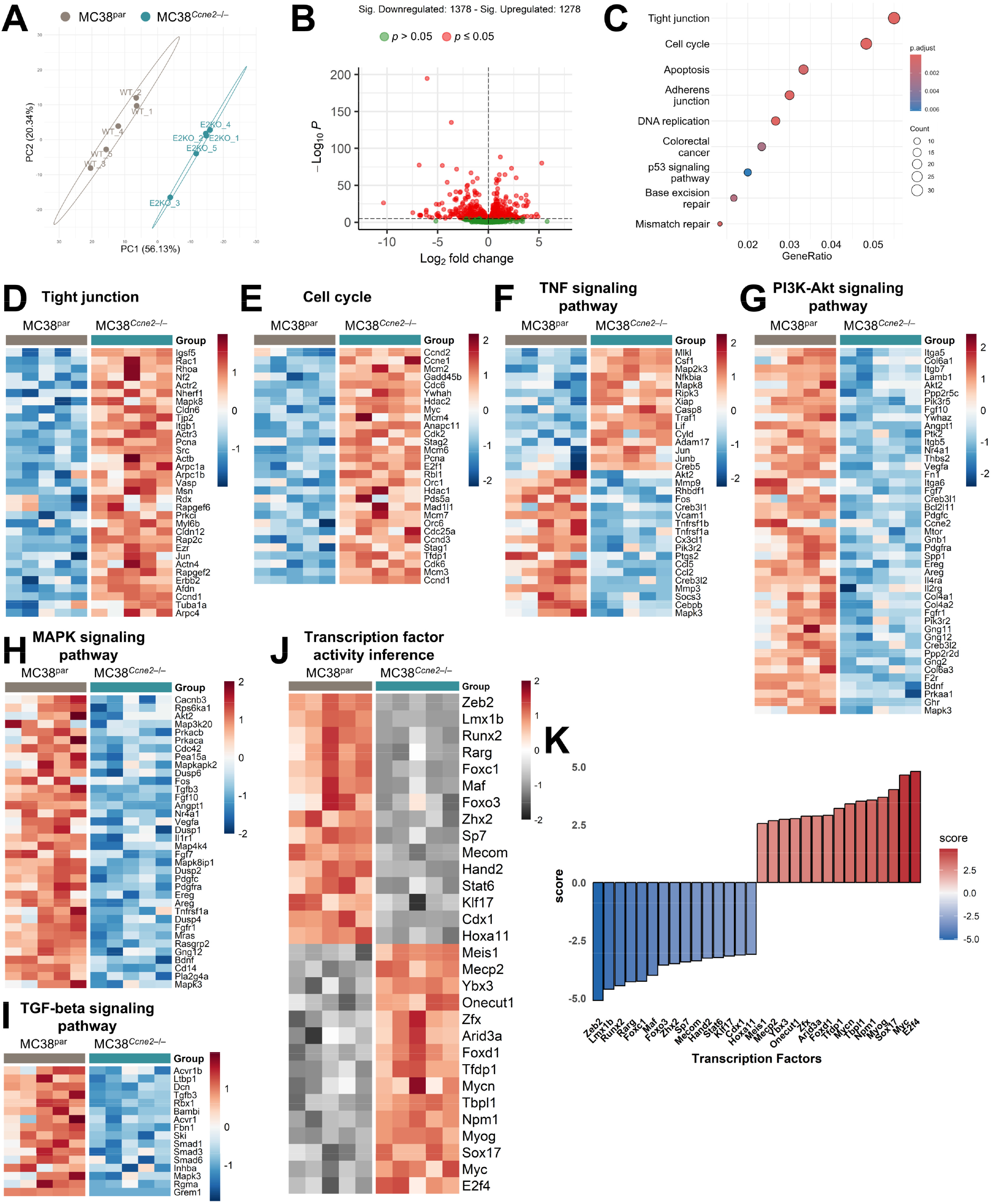
Transcriptomic profiling reveals up-regulation of TJ and cell cycle-related genes due to loss of *Ccne2* in MC38 cells. MC38 colon carcinoma cells (MC38^par^, n = 5) or *Ccne2* deleted MC38 cells (MC38*^Ccne2^*^−/−^, n = 5) were cultured for 24 h to allow cells to adhere, before replacing the medium with DMEM containing DMSO (1:1.000) for 20 h followed by RNA isolation, bulk RNA-sequencing and differential gene expression analysis. (**A**) Principal component analysis using VST normalized and batch corrected counts. (**B**) Volcano plot visualizing significantly up- and down-regulated genes (red dots: *p* < 0.05; green dots: *p* > 0.05). (**C**) Dotplot of selected pathways from over-representation analysis on KEGG pathways using significant up-regulated genes. (**D-I**) Heatmap of significant de-regulated KEGG pathways and the corresponding significant regulated genes. The color legend indicates per gene calculated z-scores. (**J**) Transcription factor activity interference analysis was performed using the decoupleR package and the CollecTRI network. The heatmap shows the 15 most significant active or inactive transcription factors in MC38*^Ccne2^*^−/−^ cells compared to MC38^par^ cells. The color legend indicates a per transcription factor scaled z-scores. (**K**) Overall transcription factor activity scores for the 15 most significant active or inactive transcription factors.

Within the TJ gene set, MC38*^Ccne2–/−^* cells showed increased expression of genes encoding cell-cell junction components and their regulators, including *Cldn6*, *Cldn12*, *Tjp2*, *Igsf5*, and the polarity/actin-remodeling regulators *Rac1*, *Rhoa*, *Cdc42*-associated effectors *Arpc1a/Arpc1b/Arpc4*, *Actb*, *Ezr*, *Msn*, *Vasp* and *Nf2* (Figure 6D). In parallel, MC38*^Ccne2–/−^* cells displayed pronounced upregulation of cell cycle genes spanning multiple phases, including origin licensing and DNA replication factors (*Mcm2*, *Mcm3*, *Mcm4*, *Mcm6*, *Mcm7*, *Orc1*, *Orc6*, *Cdc6*, *Pcna*), G1/S regulators (*Ccnd1*, *Ccnd2*, *Ccnd3*, *Cdk2*, *Cdk6*, *E2f1*, *Rb1*, *Rbl1*, *Tfdp1*, *Cdc25a*) and, notably, *Ccne1*, suggesting a compensatory transcriptional upregulation of the remaining Cyclin E paralog in response to *Ccne2* loss (Figure 6E). To complement this pathway enrichment analysis, differentially expressed cell cycle and TJ-related genes were projected onto KEGG pathway maps using Pathview (Supplementary Figure 2A-B).

Genes of the TNF signaling pathway were de-regulated in MC38*^Ccne2–/−^* cells (Figure 6F). In particular, MC38*^Ccne2–/−^* cells exhibited substantial transcriptional reprogramming of TNF-associated genes. Notably, the upregulation of *Casp8*, *Ripk3*, *Traf1*, *Nfkbia*, and *Cyld* suggests increased priming of TNF-mediated stress response and cell death pathways, whereas reduced expression of inflammatory target genes including *Ccl5*, *Ptgs2*, *Socs3*, *Mmp3*, and *Cx3cl1* points to attenuated downstream inflammatory signaling. Together, these data suggest that Cyclin E2 loss shifts TNF pathway signaling towards a state favoring stress adaptation and cell death susceptibility rather than pro-inflammatory effector responses.

In contrast, gene sets belonging to the PI3K-Akt, MAPK and TGF-beta signaling pathways were predominantly enriched among downregulated genes in MC38*^Ccne2–/−^* cells, including several growth factor and receptor tyrosine kinase components (*Pdgfra*, *Pdgfrb*, *Pdgfc*, *Fgf7*, *Fgf10*, *Fgfr1*, *Vegfa*, *Hgf*, *Mtor*), extracellular matrix/integrin genes (*Col4a1*, *Col4a2*, *Col6a1*, *Itga5*, *Itga6*, *Itgb5*, *Fn1*) and TGF-beta pathway components (*Tgfb3*, *Smad1*, *Smad3*, *Smad6*, *Acvr1b*, *Inhba*, *Grem1*) (Figure 6G-I).

To determine which transcriptional regulators might drive these expression changes, we inferred transcription factor (TF) activity using decoupleR with the CollecTRI regulon network. This analysis identified TFs with significantly divergent predicted activity between MC38^par^ and MC38*^Ccne2–/−^* cells (Figure 6J-K). Activity of the proliferation-associated TFs Myc, E2f4, Mycn, Tfdp1, Sox17, Npm1 and Tbpl1 was predicted to be increased in MC38*^Ccne2–/−^* cells, in line with the observed transcriptional upregulation of cell cycle genes upon loss of Cyclin E2. Conversely, MC38*^Ccne2–/−^* cells showed reduced predicted activity of a distinct set of TFs relative to parental cells, most prominently Zeb2, Lmx1b, Maf, Runx2 and Rarg, together with Foxc1, Foxo1, Foxo3 and Stat6. The inferred reduced activity of Zeb2, a master regulator of epithelial-to-mesenchymal transition and a transcriptional repressor of TJ and epithelial adhesion genes, is of particular interest, as its inactivation is consistent with, and may mechanistically contribute to, the concomitant upregulation of the TJ gene expression program observed in MC38*^Ccne2–/−^* cells [23, 24].

In summary, loss of *Ccne2* in MC38 cells is associated with a transcriptional program in which TJ- and cell cycle genes are over-represented among the upregulated genes, together with reduced inferred Zeb2 activity and downregulation of PI3K-Akt, MAPK and TGF-β pathway genes. Since Zeb2 represses junctional genes, its reduced activity is a candidate mechanism for the coordinated induction of the TJ program, which may contribute to the enhanced colonic barrier function observed *in vivo*.

### CDK1 overexpression is associated with increased ZO-1 and a kinase signature enriched for TJ regulators

Our *in vivo* data from *Ccne2*^−/−^ and TKO mice had demonstrated elevated *Cdk1* gene expression, particularly in colon, which was associated with reduced intestinal permeability (compare Figure 3B and 4B-C). We therefore determined whether CDK1 overexpression is sufficient to recapitulate this phenotype *in vitro*. To this end, we applied an expression construct encoding recombinant, FLAG-tagged murine CDK1 under control of a constitutively active CMV promoter and transfected it into MC38 cells. Immunoblotting confirmed robust ectopic expression of FLAG-CDK1, accompanied by a corresponding increase in total CDK1 protein, and this was associated with markedly increased expression of the TJ protein ZO-1 compared with backbone (BB)-transfected controls (Figure 7A). This finding is in agreement with the enhanced *Cdk1* and *Tjp1* gene expression observed in *Ccne2*^−/−^ and TKO colon *in vivo*.

**Figure 7:**
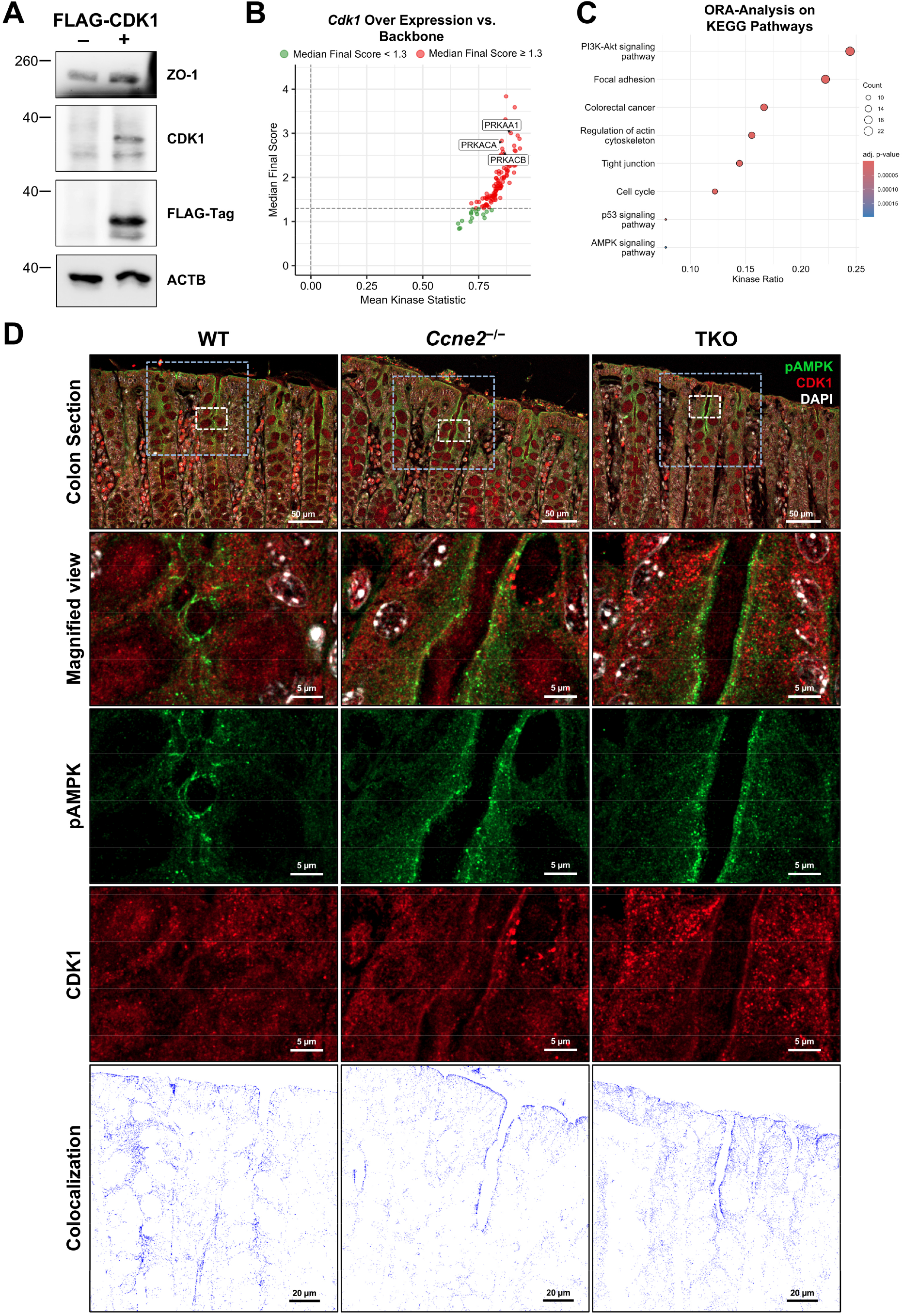
CDK1 links AMPK kinase activation to TJ protein expression and epithelial barrier integrity. (**A**) MC38 cells were transfected with an overexpression construct encoding FLAG-tagged murine CDK1 (MC38*^Cdk1^*^-OE^) or backbone control (MC38^BB^). Immunoblot analysis of ZO-1, CDK1, ectopic FLAG-tagged CDK1 and ACTB which served as loading control. (**B**) Kinomics analysis (PamDx) of serine/threonine kinase activity in MC38*^Cdk1^*^-OE^ *versus* MC38^BB^ cells. Each dot represents an individual kinase; the median final score (y-axis) reflects the specificity/confidence of the prediction; mean kinase statistic (x-axis) reflects the magnitude of activity change. Kinases exceeding the significance threshold (dashed horizontal line) are shown in red, non-significant kinases in green. The AMPK catalytic subunit PRKAA1 and the PKA catalytic subunits PRKACA and PRKACB, among the top-ranked activated kinases, are highlighted. (**C**) Over-representation analysis of significantly activated kinases against KEGG pathways. Dot size indicates the number of contributing kinases (count) and color indicates the adjusted *p*-value. (**D**) Confocal immunofluorescence microscopy of colon tissue from WT, *Ccne2*^−/−^ and TKO mice, co-stained for phospho-AMPK (pAMPK, green) and CDK1 (red) with DAPI stained nuclei (white). White dashed boxes in the upper panel correspond to enlarged views shown below; blue dashed boxes correspond to the areas selected for colocalization analysis of the pAMPK and CDK1 signals shown in the lower panels. Colocalized pixels are displayed in blue.

To identify the signaling pathways downstream of CDK1 overexpression, we profiled and compared serine/threonine kinase activity in MC38 cells with overexpression of *Cdk1* (MC38*^Cdk1^*^-OE^) and backbone transfected (MC38^BB^) cells using a chip-based kinomics platform. This assay measures real-time phosphorylation of 144 kinase-specific peptides immobilized on a 3D array and infers upstream kinase activity from the resulting phosphorylation signatures. MC38*^Cdk1^*^-OE^ cells displayed a broad shift towards overall increased kinase activity relative to controls, with most inferred active kinases exceeding a median final score > 1.2 which is considered as threshold for significant changes (Figure 7B). Among the most prominently activated kinases were the AMPK catalytic subunit PRKAA1 (AMPKα1) and the two catalytic subunits of protein kinase A, PRKACA and PRKACB, all of which ranked among the highest-scoring kinases by median final score (Figure 7B).

Over-representation analysis of the predicted significantly activated kinases against KEGG pathways revealed among others enrichment for “PI3K-Akt signaling pathway” and “Focal adhesion” as well as “Colorectal cancer”, “Regulation of actin cytoskeleton”, “Tight junction” and “Cell cycle” while “p53 signaling pathway” and “AMPK signaling pathway” were also significantly enriched, albeit with fewer contributing kinases (Figure 7C). Within the KEGG “Tight junction” pathway, the deregulated kinases comprised CDK4, the JNK family members MAPK8, MAPK9 and MAPK10, the AMPK/PKA subunits PRKAA1, PRKACA, PRKACB and PRKACG, the atypical and novel PKC isoforms PRKCE, PRKCI and PRKCZ, and the Rho-associated kinases ROCK1 and ROCK2. The latter ones collectively control TJ assembly, apical actomyosin contractility and junctional remodeling [25]. Notably, AMPK signaling emerged as a recurrent node across several of the enriched pathways. Given that AMPK is a reported CDK1 substrate during mitosis [26] and is itself a well-established positive regulator of intestinal epithelial barrier function [27], CDK1 may reinforce TJ integrity, at least in part, through activation of AMPK.

To further support the hypothesis of an association between CDK1 and AMPK signaling *in vivo*, colon sections from WT, *Ccne2*^−/−^ and TKO mice were co-stained for CDK1 and phosphorylated AMPK (pAMPK; Thr172). Confocal microscopy revealed CDK1-positive signals throughout the crypt epithelium, with a particularly intense signal in a narrow central region extending along the longitudinal crypt axis. However, CDK1 was barely found in nuclei (Figure 7D). Interestingly, pAMPK was detected within the same crypt compartment and partially overlapped with CDK1-positive areas. Colocalization analysis of higher-magnification images identified areas of spatial overlapping of CDK1 and pAMPK signals at epithelial cell borders (Figure 7D bottom panel, Supplementary Figure 3).

In addition, CDK1 also exhibited partial colocalization with ZO-1 in colonic crypt epithelial cells (Supplementary Figure 4). Together, these findings indicate a spatial association between CDK1, phosphorylated AMPK, and ZO-1, consistent with a potential role for CDK1 in the regulation of AMPK signaling at, or near, sites of cell-cell contact and epithelial barrier maintenance.

Collectively, our data support the existence of a CDK1-AMPK signaling axis associated with tight junction organization and intestinal barrier integrity both *in vitro* and *in vivo*. These findings suggest that CDK1 may contribute to epithelial barrier regulation in addition to its established role in cell cycle progression, highlighting a potential link between cell cycle-associated signaling pathways and intestinal epithelial homeostasis.

## Discussion

Using a genetic model of combined IEC-specific *Ccne1*/*Cdk2* deletion on a constitutive *Ccne2*-null background (TKO), we show that loss of the canonical Cyclin E/CDK2 module is, contrary to our initial expectation, tolerated by the intestinal epithelium and associated with enhanced barrier function. TKO and *Ccne2*^−/−^ mice display increased cell cycle activity selectively in the colon, slightly elongated crypt-villus units in the SI and reduced intestinal permeability. In *Ccne2*-deficient MC38 colon carcinoma cells, which serves as a simplified *in vitro* system, loss of Cyclin E2 was associated with a transcriptional program in which cell cycle and TJ genes were upregulated together, and CDK1 overexpression was sufficient to increase ZO-1 protein. Kinomics profiling linked CDK1 overexpression to an AMPK/PKA-related kinase network, and in colonic tissue CDK1 partially colocalized with phosphorylated AMPK and ZO-1 at epithelial cell borders. Together, these findings suggest that compensatory CDK1 activity contributes to TJ reinforcement after loss of the Cyclin E/CDK2 module or of Cyclin E2 alone, pointing to a role for CDK1 in intestinal barrier regulation beyond its canonical function in driving proliferation.

### Rationale and the unexpected direction of the phenotype

Our prior work in the liver demonstrated that concomitant inactivation of *Ccne1*, *Ccne2* and *Cdk2* in hepatocytes is compatible with normal organ homeostasis under basal, non-regenerative conditions but severely compromises compensatory hepatocyte proliferation after two-thirds partial hepatectomy [8]. Because the intestinal epithelium undergoes continuous self-renewal even under basal conditions, with a turnover time of only a few days, we initially hypothesized that combined loss of the entire Cyclin E/CDK2 module would be poorly tolerated in the gut and might phenocopy the regenerative failure observed in the liver. However, the data presented here contradict this hypothesis. Rather than impairing epithelial renewal, loss of *Ccne2* alone, or of the complete Cyclin E1/E2/CDK2 module, is associated with preserved or even enhanced cell cycle activity restricted to the colon, normal tissue architecture, and remarkably reduced intestinal permeability under basal conditions. This apparent contradiction is best reconciled by the existence of redundant, compensatory cell cycle entry pathways in the gut that are not available or not efficiently engaged in hepatocytes. In the intestinal crypt, a constitutively active stem-cell compartment together with a permissive proliferative niche may allow for rapid substitution of the canonical G1/S Cyclin E/CDK2 checkpoint by other cyclin-CDK complexes, most notably Cyclin B1/CDK1 that we find transcriptionally induced in TKO and *Ccne2*^−/−^ colon. This is consistent with the known capacity of CDK1 to compensate for the loss of interphase CDKs in several tissue contexts [28, 29], and it illustrates that the requirement for individual cell cycle regulators is highly tissue- and context-dependent rather than a fixed, universally applicable rule of the canonical cell cycle machinery.

### Reduced Mendelian ratios and litter size in TKO breeding

A notable and unexpected observation was the lower-than-expected proportion of TKO offspring from respective breeding pairs, together with smaller litter sizes compared with WT breedings. Because *Ccne2*^−/−^ male mice are known to be largely infertile due to a meiotic defect [18], our breeding scheme relied on heterozygous males, and germline transmission of the floxed *Cdk2* and *Ccne1* alleles should, in principle, be restricted to Villin1-Cre-negative cells. However, a recent report demonstrated that the Villin1 promoter in several widely used Cre driver lines exhibits substantial “leaky” recombinase activity outside the intestinal epithelium, including within subsets of seminiferous tubules in the testis [30]. This raises the plausible possibility that Cre-mediated excision of the floxed *Cdk2* and *Ccne1* alleles also occurred, at least partially, in the male germline of our breeding animals, independent of intestinal Villin1 promoter activity. Given the established, non-redundant requirement of *Cdk2* for meiotic progression [20], unintended germline deletion of *Cdk2* could readily explain both reduced male fertility and a biased transmission ratio of the fully recombined TKO allele combination, without invoking any developmental function of Cyclin E/CDK2 in the gut itself. Alternatively, non-mutually exclusive explanation is that Cyclin E/CDK2 activity is required during early post-implantation development of the gut endoderm or during the establishment of the intestinal stem-cell compartment, a period that precedes robust Villin1-Cre activity and during which even partial mosaic recombination in extra-intestinal or extra-embryonic tissues could compromise embryonic viability. However, as we were able to clearly demonstrate that TKO mice are generally viable and do not exhibit any pronounced developmental phenotype after birth, the aspects mentioned above were not investigated further.

### A model linking Cyclin E2 loss to compensatory cell cycle activation and TJ reinforcement

We propose that *Ccne2* loss in the colonic epithelium triggers compensatory cell cycle remodeling, including increased *Ccnb1* and *Cdk1* expression, which promotes a TJ program and strengthens barrier function (compare Figure 4A). This idea was largely supported by our MC38 cell culture model: In proliferating cells, *Tjp1* expression closely correlated with *Cdk1*, but not with *Ccna2* or *Ccnb1*. *Ccne2*-deficient MC38 cells showed enhanced transcriptional activation of the general cell cycle machinery, which was associated with increased ZO-1 protein expression and coordinated transcriptional induction of further TJ-related genes. Moreover, CDK1 overexpression alone was sufficient to increase ZO-1 protein indicating that CDK1 can directly promote TJ regulation. This is consistent with recent evidence that ZO-1 is a CDK1 substrate at epithelial cell-cell contacts [31]. Thus, compensatory cell cycle rewiring after *Ccne2* loss may reinforce the epithelial barrier through both increased junctional gene expression and direct CDK1-dependent regulation of ZO-1.

### CDK1 as a direct regulator of TJs

The concept that CDK1, a kinase traditionally regarded as a purely mitotic, chromatin- and spindle-associated enzyme, might also control epithelial barrier integrity was entirely unexpected at the beginning of this study. Our kinomics profiling of CDK1-overexpressing MC38 cells revealed broad activation of serine/threonine kinases enriched for the KEGG TJ pathway (CDK4, MAPK8/9/10, PRKAA1, PRKACA/B/G, PRKCE/I/Z, ROCK1/2) compared to control MC38 cells, and our immunofluorescence and confocal imaging of TKO colon directly visualized a CDK1 pool concentrated in a discrete, non-nuclear, junction-proximal compartment that partially co-localized with active phospho-AMPK and with ZO-1 at the lateral epithelial membrane. These findings gain strong, and considerably more detailed independent support from the very recent report by Dantas et al. [31]. This study used a confluent epithelial monolayer formed by Madin-darby canine kidney (MDCK) cells, which is a model largely unrelated to intestinal epithelium, and identified CDK1 as a regulator of TJ proteins and epithelial barrier integrity. In particular, phosphoproteomics established ZO-1 as a direct CDK1 target, thereby supporting the conclusion that CDK1 can regulate intestinal TJs through direct phosphorylation of ZO-1. Together with the observation that this junctional CDK1 activity persists throughout G1-, S- and G2-phase rather than being restricted to mitosis, the data from Dantas et al. provide an independent, mechanistically direct explanation for our own findings: CDK1 might not act solely through downstream kinase cascades such as AMPK to reinforce TJs, but can phosphorylate ZO-1 and its junctional partners directly. This raises the intriguing possibility that the CDK1-AMPK association observed at the lateral membrane *in vivo* reflects a parallel or convergent arm of the same junctional signaling hub, particularly given that AMPK itself has been shown to phosphorylate Afadin to promote ZO-1 recruitment to the plasma membrane [27]. Whether CDK1 and AMPK act on overlapping or distinct residues of ZO-1 and Afadin remains to be determined. Irrespective of this open question, the convergence between two independently generated, methodologically orthogonal datasets supports the conclusion that CDK1 is a *bona fide*, spatially restricted regulator of epithelial junctional architecture and barrier function in addition to its canonical role in driving mitotic entry.

### Reduced ZEB2 activity as a candidate transcriptional driver of the TJ program

Our transcription factor activity inference analysis predicted reduced ZEB2 activity in MC38*^Ccne2^*^−/−^ cells. ZEB2 is a transcription factor that represses a broad program of epithelial cell-cell junction genes, including TJ, adherens junction, desmosomal and gap junction components [32]. In skin epithelium, conditional ZEB2 overexpression is sufficient to compromise epidermal barrier integrity through direct transcriptional repression of occludin and other TJ genes, establishing a causal, tissue-relevant link between ZEB2 activity and barrier function that mirrors what we observe in reverse upon ZEB2 loss of activity in MC38*^Ccne2^*^−/−^ cells [23]. In the intestine, ZEB2 has been shown to drive an invasive, microbiota-dependent colon carcinoma phenotype, through disruption of epithelial cell-cell adhesion programs [24]. Our finding that *Ccne2* loss predicts reduced ZEB2 activity therefore provides a plausible explanation for the coordinated activation of the TJ gene expression program observed by RNA-seq analysis. However, ZEB2 activity was not measured directly so far. Thus, reduced ZEB2 activity should be considered a hypothesis-generating candidate mechanism explaining our current findings that requires experimental validation in future studies.

### Limitations of the study and open questions

Although our findings establish an unexpected link between compensatory cell cycle control and epithelial barrier regulation, several limitations should be acknowledged. The genetic strategy employed does not fully distinguish epithelial-intrinsic effects from potential systemic consequences of constitutive *Ccne2*-deficiency. Moreover, parts of the mechanistic analysis rely on a transformed epithelial cell model and therefore warrant validation in additional physiological systems. While the data collectively support a CDK1-associated signaling network that coordinates proliferation and tight junction organization, the precise molecular architecture of this pathway remains incompletely defined. Finally, the study was performed under steady-state conditions, and the relevance of this adaptive response in settings of intestinal injury or inflammation remains to be determined.

### Therapeutic outlook

The notion that transient perturbation of key cell cycle regulators could therapeutically reinforce the intestinal barrier appears paradoxical, considering their well-established role in oncogenic proliferation. However, our findings suggest that transient, tightly controlled activation of CDK1 could be explored as a strategy to reinforce the intestinal barrier during acute mucosal injury. For instance, short term WEE1 inhibition by the clinically investigated inhibitor adavosertib, which increases CDK1 activity by reducing inhibitory Thr15 phosphorylation [33], could be one appropriate approach. Given the proliferative and genotoxic risks of sustained cell cycle activation, such interventions would require low-dose, pulsed administration and preferential restriction to the colon through appropriate delivery routes and formulations. Comparative preclinical studies should define which application mode offers optimal balance between epithelial barrier reinforcement, tissue distribution and safety.

### Concluding remarks

Collectively, this study identifies an unexpected, barrier-protective consequence of Cyclin E/Cdk2 loss in the murine colon and proposes a model that connects the intestinal epithelial cell cycle machinery to barrier by proceeding from compensatory *Cdk1* induction, through direct and indirect upregulation of TJ-related genes and proteins to eventually reinforced TJ assembly. Beyond adding CDK1 to the list of cell cycle regulators with functions in epithelial junctional biology, our findings raise the broader possibility that transient, pharmacologically induced modulation of this axis could one day be explored as a novel strategy to reinforce a compromised intestinal barrier in acute mucosal disease.

## Materials and Methods

### Animal maintenance and experimentation

All animal experiments were approved by the State Office for Nature, Environment and Consumer Protection of North Rhine-Westphalia (LANUV, Recklinghausen, Germany, AZ-84-02.04.2017.A256) and performed in accordance with German animal welfare legislation. Mice were housed at the Institute of Laboratory Animal Science, Uniklinik RWTH Aachen, under controlled environmental conditions with a 12 h light/dark cycle and *ad libitum* access to standard chow diet and water. All mouse lines were maintained on a C57BL/6J background. Constitutive *Ccne2*^−/−^ mice [18] and mice carrying floxed *Ccne1* alleles [19] were kindly provided by P. Sicinski (Dana-Farber Cancer Institute, Boston, USA); mice carrying floxed *Cdk2* alleles [20] were provided by M. Barbacid (Spanish National Cancer Research Centre, Madrid, Spain). The generation of *Ccne1*^f/f^ *Cdk2*^f/f^ mice on a *Ccne2*^−/−^ background has basically been described previously [8]. To delete *Ccne1* and *Cdk2* in intestinal epithelial cells (IECs) of the small and large intestine, *Ccne1*^f/f^ *Ccne2*^−/−^ *Cdk2*^f/f^ mice were crossed with transgenic mice expressing Cre recombinase under the control of the *Villin 1* promoter [17]. The resulting *Ccne1*^ΔIEC^ *Ccne2*^−/−^ *Cdk2*^ΔIEC^ mice (triple knockout, TKO) lack the complete Cyclin E/CDK2 complex in IECs. Cre-negative *Ccne1*^f/f^ *Ccne2*^−/−^ *Cdk2*^f/f^ mice, which lack only Cyclin E2, and Cre-negative *Ccne1*^f/f^ *Ccne2*^+/+^ *Cdk2*^f/f^ mice served as *Ccne2*^−/−^ and wildtype (WT) controls, respectively. Because homozygous *Ccne2*^−/−^ males are sterile, heterozygous *Ccne2* males were used for breeding.

For *in vivo* intestinal permeability assays, mice of both genders were fasted for 3 h and received 4 kDa FITC-dextran (Sigma-Aldrich) at a dose of 440 µg/g body weight by oral gavage. After 4 h, mice were euthanized by cervical dislocation and blood was collected from the *vena cava*. Samples were protected from light throughout. Serum was obtained by centrifugation (10,000 × *g*, 5 min, room temperature), diluted 1:3 in DPBS, and fluorescence was measured (excitation 485 nm, emission 530 nm) using a Cytation 3 Cell Imaging Multi-Mode Reader (BioTek Instruments, Winooski, VT, USA). FITC-dextran concentrations were calculated from a standard curve of serially diluted FITC-dextran.

For the collection of tissues, mice were euthanized by cervical dislocation, and body weight was recorded. Livers were weighed and portions were fixed in 4 % formaldehyde for 24 h at 4 °C with subsequent dehydration and paraffin embedding or snap-frozen in liquid nitrogen and stored at −80 °C. After removal of the mesentery, the lengths of small intestine and colon were measured. Both were flushed with DPBS (18G gavage needle) and opened longitudinally; the small intestine was divided into proximal, medial and distal segments. Intestinal segments were snap-frozen for RNA and protein isolation or fixed in methacarn for 24 h at 4 °C with subsequent dehydration and paraffin embedding for immunohistochemistry.

### Immunohistochemistry, immunofluorescence staining and microscopy

Slicing and hematoxylin and eosin staining of fixed paraffin embedded liver and intestinal tissues was performed by the Immunohistochemistry Facility of the Interdisciplinary Center for Clinical Research (IZKF) of the RWTH Aachen following standard procedures. Whole slide images were acquired using a Zeiss Axio Observer.Z1 microscope (Carl Zeiss Microscopy, Jena, Germany) equipped with a motorized stage. Shading correction of individual images and tile stitching was performed using Zen Pro (v2.6) software.

Immunohistochemistry staining on paraffin embedded sections of liver and colon tissues was performed using a monoclonal CD45 antibody (Clone D3F8Q, Cell Signaling Technology) or a Ki67 antibody (Clone SP6, abcam) together with an anti-rabbit IgG HRP-linked secondary antibody (#7074, Cell Signaling Technology) following standard staining procedures. CD45 positive area fractions in liver tissues were quantified using ImageJ (v1.54.h) software.

Immunofluorescence staining of paraffin embedded colon sections was performed as previously described [34]. In brief, paraffin sections were rehydrated using xylol, ethanol and distilled water followed by incubation in IHC Antigen Retrieval Solution with low pH (Invitrogen) and subsequently with Image-iT FX Signal Enhancer (Invitrogen), blocking with 5 % goat serum and 0.1 % Triton X-100 in DPBS, supplemented with 10 % Goat F(ab) Anti-Mouse IgG H&L (abcam, ab6668) prior to incubation with primary antibodies for CDK1 (Invitrogen, MA5-11472), ZO-1 (abcam, ab96587) or phospho-AMPK (Thr172) (Cell Signaling, 2535S) at 4 °C overnight. After removal of primary antibodies, slices were incubated with fluorescence labeled secondary antibodies for 2 h at room temperature followed by DAPI staining before mounting with ProLong Diamond mounting medium (Invitrogen).

Fluorescence images were acquired using an inverted Zeiss LSM 980 confocal laser scanning microscope equipped with a Plan-Apochromat 10x/0.45 M27 and a Plan-Apochromat 40x/1.3 NA oil-immersion DIC (UV) VIS-IR M27 objective. Signals were acquired sequentially in three tracks to minimize spectral crosstalk between fluorophores. Alexa Fluor 647 was excited using the 639 nm laser line at 0.6 % laser power, Alexa Fluor 488 using the 488 nm laser line at 1.2 %, and DAPI using the 405 nm laser line at 0.2 %. Fluorescence was detected using the LSM 980/Airyscan detection system with GaAsP-PMT detection. Airyscan acquisition was performed using the FastAiryScan Sheppard-sum mode.

Colocalization was analyzed in Zen (v3.0, Zeiss) using the colocalization software module by applying thresholds on both Alexa Fluor 488 and 647 channels, to identify pixels which are double positive for both fluorescence signals as shown in the respective colocalization scatter plots. Double positive pixels were highlighted to visualize areas of colocalization.

### Cell culture

MC38 murine colon carcinoma cells (Sigma-Aldrich, St. Louis, MO, USA) were thawed and expanded in MC38 expansion medium consisting of high-glucose Dulbecco’s Modified Eagle’s Medium (DMEM, PAN-Biotech GmbH, Aidenbach, Germany) supplemented with 10 % fetal bovine serum (FBS, PAN-Biotech), 2 mM L-glutamine (Sigma-Aldrich), 1x non-essential amino acids (Sigma-Aldrich), 1 mM sodium pyruvate (Sigma-Aldrich), and 50 µg/mL gentamycin sulfate (Sigma-Aldrich) in T75 flasks (VWR International Europe, Leuven, Belgium). Cells were maintained at 37 °C in a humidified incubator containing 5 % CO_2_. Upon reaching approximately 80-90 % confluency, cells were detached using trypsin-EDTA (PAN-Biotech) and routinely passaged at a 1:100 split ratio in fresh MC38 expansion medium. Parental and genetically modified MC38 cell lines were cultured under identical conditions unless otherwise stated. All cell lines were routinely tested for mycoplasma contamination and were confirmed to be negative.

### Generation of MC38*^Ccne2^*^−/−^ cells

To recapitulate the genetic background of our animal models *in vitro*, we generated a *Ccne2* knockout in MC38 cells using a CRISPR/Cas9-based approach. To address this, a guide RNA (gRNA) was created using a murine *Ccne2*-specific 23 bp long CRISPR RNA (crRNA) against *Ccne2*, and an ATTO550-labeled trans-activating crRNA (tracrRNA). The crRNA target sequence was *5′-GAGTTTGCTTACGTCACTGA-3′*. Target-specific crRNA, tracrRNA and Alt-R™ S.p. Cas9 Nuclease V3 were obtained from Integrated DNA Technologies (IDT, Leuwen, Belgium). Parental MC38 cells were transfected with preassembled gRNA-Cas9 ribonucleoprotein complexes consisting of gRNA and Alt-R™ S.p. Cas9 Nuclease V3 using Lipofectamine™ CRISPRMAX™ Cas9 Transfection Reagent (Thermo Fisher Scientific) according to the manufacturer’s instructions. After 48 h, transfected single cell clones were sorted based on the fluorescence signal of ATTO550-labeled crRNA using a BD FACSAria™ Fusion (Becton Dickinson, New Jersey, USA). Single-cell-derived clones were expanded in culture medium and maintained until they reached full confluence in a 6-well plate format before further analysis. Two step verification for targeted deletion of *Ccne2* was achieved *via* melting curve analysis of locus-specific qPCR fragments followed by Sanger Sequencing of selected clones provided by Eurofins Genomics (Ebersberg, Germany). A clone with a suitable frameshift mutation and a predicted loss of function was selected for subsequent downstream analyses. The prediction of the protein structure of the truncated, mutated Cyclin E2 protein was generated using AlphaFold 2 [22]. Molecular graphics and analyses were performed with UCSF ChimeraX [35].

### Plasmid generation and transfection

To investigate the effect of increased *Cdk1* expression in MC38 cells, we designed a plasmid encoding murine CDK1 fused to an N-terminal 3xFLAG-tag and co-expressing EGFP fused to a nuclear localization signal referred to as pCdk1-OE. The pCdk1-OE vector used to overexpress *Cdk1* in our study as well as the respective control plasmid (pBB) were constructed by VectorBuilder (Neu-Isenburg, Germany). The vector ID is [VB250415-1084ekt] for pCdk1-OE and [VB250415-1197nqd] for pBB, which can be used to retrieve detailed information about the vector on www.vectorbuilder.com. Transient plasmid transfection into MC38 cells was performed using Lipofectamine™ 3000 transfection reagent (Thermo Fisher Scientific) according to manufacturer’s instructions. After transfection, cells were maintained under standard culture conditions for 24 h before further analysis.

### RNA isolation and quantitative Real-Time PCR (qPCR)

Total RNA was extracted from gut and liver tissue samples and cell pellets using QIAzol Lysis Reagent (Qiagen, Hilden, Germany). Complementary DNA (cDNA) was synthesized by reverse transcription utilizing the Omniscript RT Kit (Qiagen). Relative mRNA expression levels were quantified by real-time quantitative PCR (qPCR) using QuantStudio 5 and QuantStudio 6 Flex Real-Time PCR instruments (Applied Biosystems, Foster City, CA, USA) in combination with Fast SYBR Green PCR Master Mix (Invitrogen, Carlsbad, CA, USA). Gene expression was analyzed using the 2^−ΔΔCt^-method [36] with *Gapdh* as reference gene for normalization and is presented as fold changes relative to the respective control group, as specified in the figure legends. Alternatively, correlation analysis was performed using *Gapdh*-normalized gene expression values (-ΔCt). Pairwise Pearson correlation coefficients were calculated and visualized using the ggpairs() function of the GGally package (v2.4.0) in R (v4.5.1). All primer sequences applied in this study are provided in Table 1.

**Table 1:**
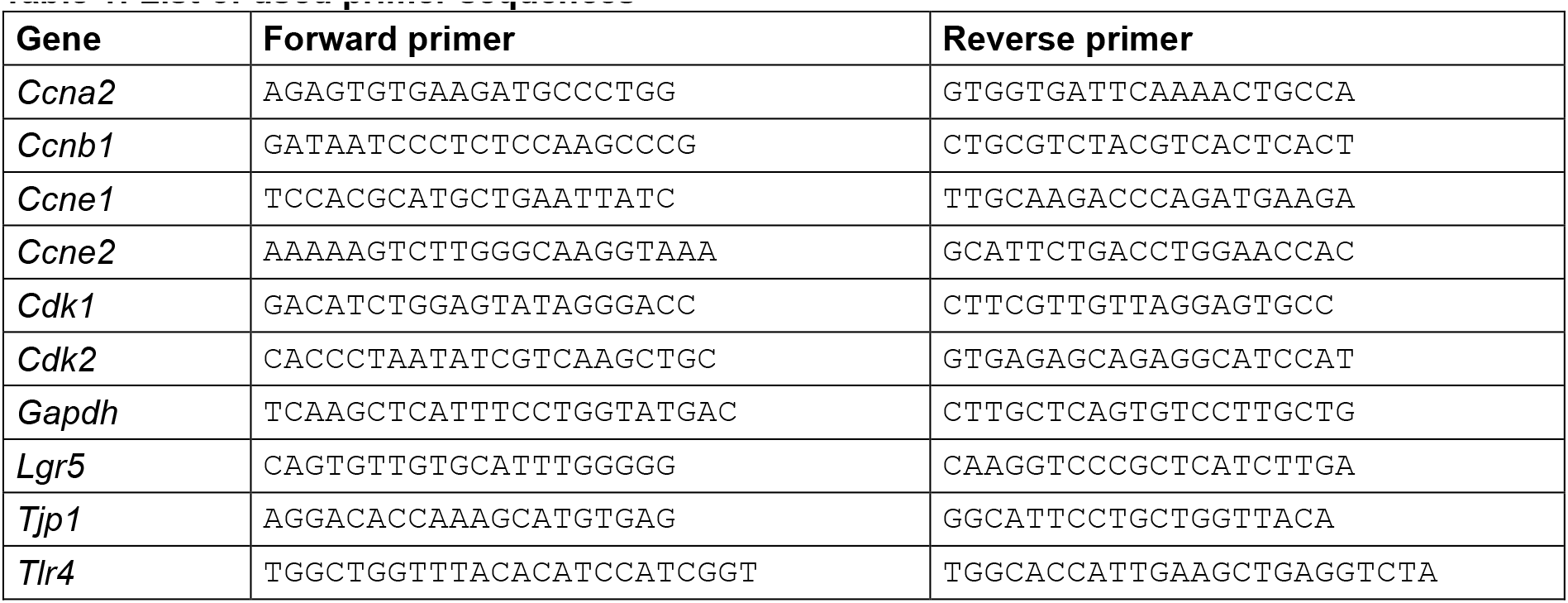
List of used primer sequences.

**Table 2:** List of primary antibodies used for immunoblot analysis.

| Protein | Primary antibody | Species | Provider |
| --- | --- | --- | --- |
| ACTB | beta-Actin (13E5) Monoclonal Antibody | rabbit | Cell Signaling |
| CCNE2 | Anti-Cyclin E2 antibody [E142] | rabbit | abcam |
| CDK1 | CDK1 Monoclonal Antibody (A17.1.1) | mouse | Invitrogen |
| ZO-1 | Anti-ZO1 tight junction protein antibody | rabbit | abcam |
| pCDK substrate | pCDK Substrate (pTPXK) (D9V5N) | rabbit | Cell Signaling |
| FLAG | Monoclonal ANTI-FLAG M2 antibody | mouse | Merck |

### Immunoblot analysis

Tissues were lysed in ice-cold NP40-Lysis Buffer using a PTFE tissue grinder (VWR, Darmstadt, Germany). Western blots were carried out according to standard procedures [10]. Membranes (Whatman® Protran®, Maidstone, UK) were probed with primary antibodies as listed in Table 1. As secondary antibodies, horseradish peroxidase (HRP)-conjugated anti-rabbit IgG (Cell signaling) and anti-mouse-HRP (m-IgGκ BP-HRP, Santa Cruz) were used. Uncropped Western blots can be found in Supplementary Figure 5.

### EdU incorporation and flow cytometric cell cycle analysis

Parental and genetically modified MC38 cells were seeded in 6-well plates and analyzed 24 h after seeding. To assess S-phase activity, cell culture medium was supplemented with 10 µM 5-ethynyl-2′-deoxyuridine (EdU) for 30 min before harvest. Cells were detached using trypsin-EDTA, resuspended in culture medium, and transferred to flow cytometry tubes. Cells were collected by centrifugation, washed with 1 % bovine serum albumin in Dulbecco’s phosphate-buffered saline (DPBS, PAN-Biotech). Cells were fixed in permeabilized using the Foxp3/Transcription Factor Staining Buffer Set (Thermo Fisher Scientific, Waltham, MA, USA) as previously described [37]. Incorporated EdU was detected using the Click-iT Plus EdU Alexa Fluor 488 Flow Cytometry Assay Kit (Thermo Fisher Scientific) following manufacturer’s instructions followed by additional antibody staining of phosphorylated histone H3 (Ser10, Cell Signaling Technology, Danvers, MA, USA). After washing cells were resuspended in DPBS containing DAPI at a final concentration of 1:500 and incubated for 10 min. Samples were measured using a BD LSRFortessa™ Cell Analyzer (BD, Franklin Lakes, New Jersey, USA) and were analyzed using FlowJo software (v10, BD). The applied gating strategy is shown in Supplementary Figure 6.

### RNA sequencing and transcriptome analysis of MC38-derived cell lines

MC38^par^ and MC38*^Ccne2^*^−/−^ cell lines were seeded and cultured for 24 h prior to treatment with DMSO (1:1000). After an additional 20 h incubation, total RNA was isolated for RNA sequencing. Sequencing libraries were prepared using the QuantSeq 3’mRNA-Seq Library Prep Kit FWD for Illumina (Lexogen, Vienna, Austria) following manufacturer’s protocols. Libraries were sequenced on a NextSeq 500 system using the high output kit v2.5 (75 cycles) (Illumina, San Diego, USA). Raw reads were processed using the nf-core/rnaseq pipeline (v3.17.0) [38] including read alignment to the GRCm39 reference genome using STAR (v2.7.11b) [39] and read quantification with Salmon (v1.10.3) [40]. Count normalization and differential gene expression analysis was performed in R (v4.5.1) using DESeq2 (v1.48.2) [41]. Genes with adjusted *p* < 0.05 after Benjamini-Hochberg correction were considered differentially expressed. For overrepresentation analysis of differentially up- or down-regulated genes on KEGG pathways, clusterProfiler (v4.16.0) [42] was used. Results of overrepresentation analysis were visualized using Pathview 1.52.0 [43]. Transcription factor activity inference analysis was performed using the decoupleR package (v2.14.9) [44] and the CollecTRI network.

### Kinase activity profiling

MC38 cells were lysed for 10 min on ice using M-PER Mammalian Protein Extraction Reagent (Thermo Scientific, cat no. 78503) containing Halt Phosphatase Inhibitor Cocktail and Halt Protease Inhibitor Cocktail EDTA-free (1:100 each; Thermo Scientific, cat no. 78420 and 87785). The lysates were centrifuged at 15,000 × g for 15 min at 4 °C. In accordance with the manufacturer’s instructions, protein quantification was determined using the PierceTM BCA Protein Assay Kit (Thermo Scientific).

Serine-Threonine kinase profiles of MC38-derived cell lysates (n = 5 per genotype, 1.0 μg of protein) were determined using the PamChip® Ser/Thr Kinase assay (STK; PamDx International, ’s-Hertogenbosch, The Netherlands) as described recently [45] with minor modifications. Each STK array contains 144 individual phospho-site(s) that are peptide sequences derived from substrates for Ser/Thr kinases. The peptides on the chip builds a 15-amino acid sequence representing a putative endogenous phosphorylation site which functions as a serine-threonine kinase substrate. A FITC-conjugated antibody was applied to visualize phosphorylation after 1 h of incubation at 30 °C. Imaging was performed using an CCD camera (PamStation®12), and spot intensities were measured at each time point and adjusted for local background using the BioNavigator software version (v6.3, PamDx). Upstream Kinase Analysis [46], a functional scoring method, was used to rank kinases based on combined specificity scores (based on peptides linked to a kinase, derived from 6 databases) and sensitivity scores (based on treatment-control differences). Data analysis was performed in R (v4.5.1) using the EnhancedVolcano package (v1.26.0) to visualize significant deregulated inferred kinase activity based on a significance threshold defined by a median final score > 1.2. Over-representation of differentially active kinases on KEGG pathways was analyzed with the clusterProfiler package (v4.16.0).

### Isolation and culture of murine small-intestinal organoids

Murine small-intestinal organoids were established from 8-week-old mice. The small intestine was removed, flushed with ice-cold phosphate-buffered saline (DPBS; PAN-Biotech), opened longitudinally, and cleared of villi by gentle scraping. After repeated washing, tissue fragments were sequentially incubated in 1 mM and 5 mM EDTA (AppliChem, Darmstadt, Germany) in DPBS for 30 and 60 min, respectively, at 4 °C with continuous rotation. Crypts were released by vigorous mechanical agitation, passed through a 70 µm cell strainer, and collected by centrifugation at 300 × *g* for 5 min at 4 °C. Approximately 2,500 crypts were embedded in 20 µl Matrigel Matrix (Corning, Corning, NY, USA) per well of a prewarmed 48-well plate.

After polymerization at 37 °C, Matrigel domes were overlaid with ENR medium consisting of Advanced DMEM/F12 supplemented with 1 % GlutaMAX, 10 mM HEPES, and 1 % penicillin–streptomycin (all from Thermo Fisher Scientific), 1x B27 and 1x N2 supplements and recombinant murine EGF (50 ng/mL; Thermo Fisher Scientific), *N*-acetyl-L-cysteine (1.25 mM; Sigma-Aldrich), recombinant murine Noggin (100 ng/mL; PeproTech, Hamburg, Germany), and recombinant human R-spondin 1 (250 ng/mL; R&D Systems, Minneapolis, MN, USA). Organoids were maintained at 37 °C in a humidified atmosphere containing 5 % CO_2_, with medium replacement every 3 days. Organoids were passaged every 7 to 10 days depending on size.

### Statistical analysis

Statistical analysis was performed using Graph Pad Prism 11 (Graphpad Software, Inc., USA). For comparison of two groups, an unpaired, two-tailed *t* test was used. Comparison of groups greater than or equal to three was performed using ordinary one-way ANOVA followed by appropriate multiple comparison tests as indicated in the respective figure legends. The analysis of groups differing in two variables (*e.g.* time point and genotype) was done by two-way ANOVA followed by Šídák’s multiple comparisons test. All data are presented as mean ± SD. Significances were defined as *: *p* < 0.05; **: *p* < 0.01; ***: *p* < 0.001; ****: *p* < 0.0001.

## Data availability

The RNA-sequencing data and other data supporting the findings of this study are available from the corresponding author upon reasonable request.

## Acknowledgements

This work was supported by the German Research Foundation (DFG/CRC 1382 “Gut-Liver Axis”, Project No. 403224013, to CL (project A02) and to SLM (“rising star initiative for emerging PIs”), and to E.P.C.v.d.V (INST 222/1598-1 FUGG; Projekt No. 566187929 and Project No. 569469700)), and by the Corona Foundation (S199/10084/2021) to E.P.C.v.d.V.. YAN was funded by MICIU/AEI (grants PID2023-150260OB-I00 and 10.13039/501100011033) and by HORIZON-HLTH-2022-STAYHLTH-02 (agreement No 101095679).

This work was supported by the Flow Cytometry Facility (RRID:SCR_028757), the Confocal-Microscopy-Facility, the Immunohistochemistry Facility and the Genomics Facility of the Interdisciplinary Center for Clinical Research (IZKF) within the Faculty of Medicine at RWTH Aachen University.

## Abbreviations

ACTB: β-actin
AMPK: AMP-activated protein kinase
BB: backbone control
CDK: cyclin-dependent kinase
DAPI: 4′,6-diamidino-2-phenylindole
EdU: 5-ethynyl-2′-deoxyuridine
EGF: epidermal growth factor
EGFP: enhanced green fluorescent protein
FITC: fluorescein isothiocyanate
gRNA: guide RNA
H&E: hematoxylin and eosin
IEC: intestinal epithelial cell
ISC: intestinal stem cells
MC38par: parental MC38 cells
MC38Ccne2–/–: MC38 cells with Ccne2 knockout
MC38Cdk1-OE: MC38 cells overexpressing Cdk1
PCNA: proliferating cell nuclear antigen
PI3K: phosphoinositide 3-kinase
PKA: protein kinase A
SI: small intestine
STK: serine/threonine kinase
TF: transcription factor
TGF-β: transforming growth factor beta
TJ: tight junction
TKO: Ccne1ΔIEC Ccne2–/– Cdk2ΔIEC triple-knockout mice
TNF: tumor necrosis factor
WT: wildtype
ZEB2: zinc finger E-box-binding homeobox 2
ZO-1: zonula occludens-1

---

**Supplementary Figure 1:**
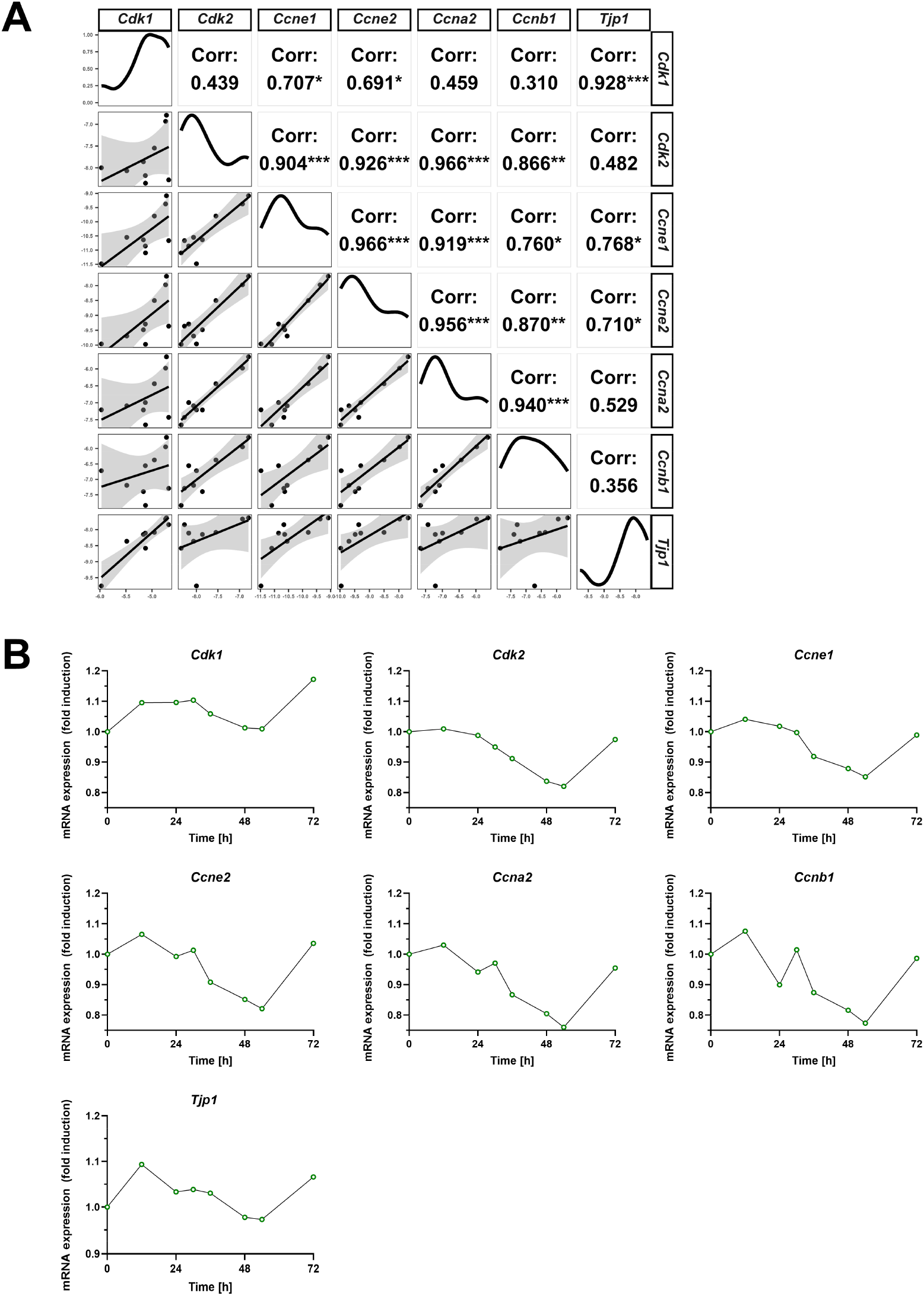
Continuously proliferating MC38 cells were collected regularly over 72 hours and relative mRNA expression of *Cdk1*, *Cdk2*, *Ccne1*, *Ccna2*, *Ccnb1* and *Tjp1* was determined by qPCR (n = 1). (**A**) Expression was calculated as -ΔCt using the housekeeping gene *Gapdh* for normalization followed by gene-wise correlation analysis. Correlation plots are shown in the lower left corner and Pearson correlation statistics including significance are shown in the upper right corner. (**B**) Time-coures analysis of the indicated genes. Expression levels were normalized to *Gapdh* expression and calculated as fold induction compared to the 0 h time point. Pearson correlation statistics were calculated using the GGally package in R (**A**). Significances were defined as *: *p* < 0.05; **: *p* < 0.01; ***: *p* < 0.001

**Supplementary Figure 2:**
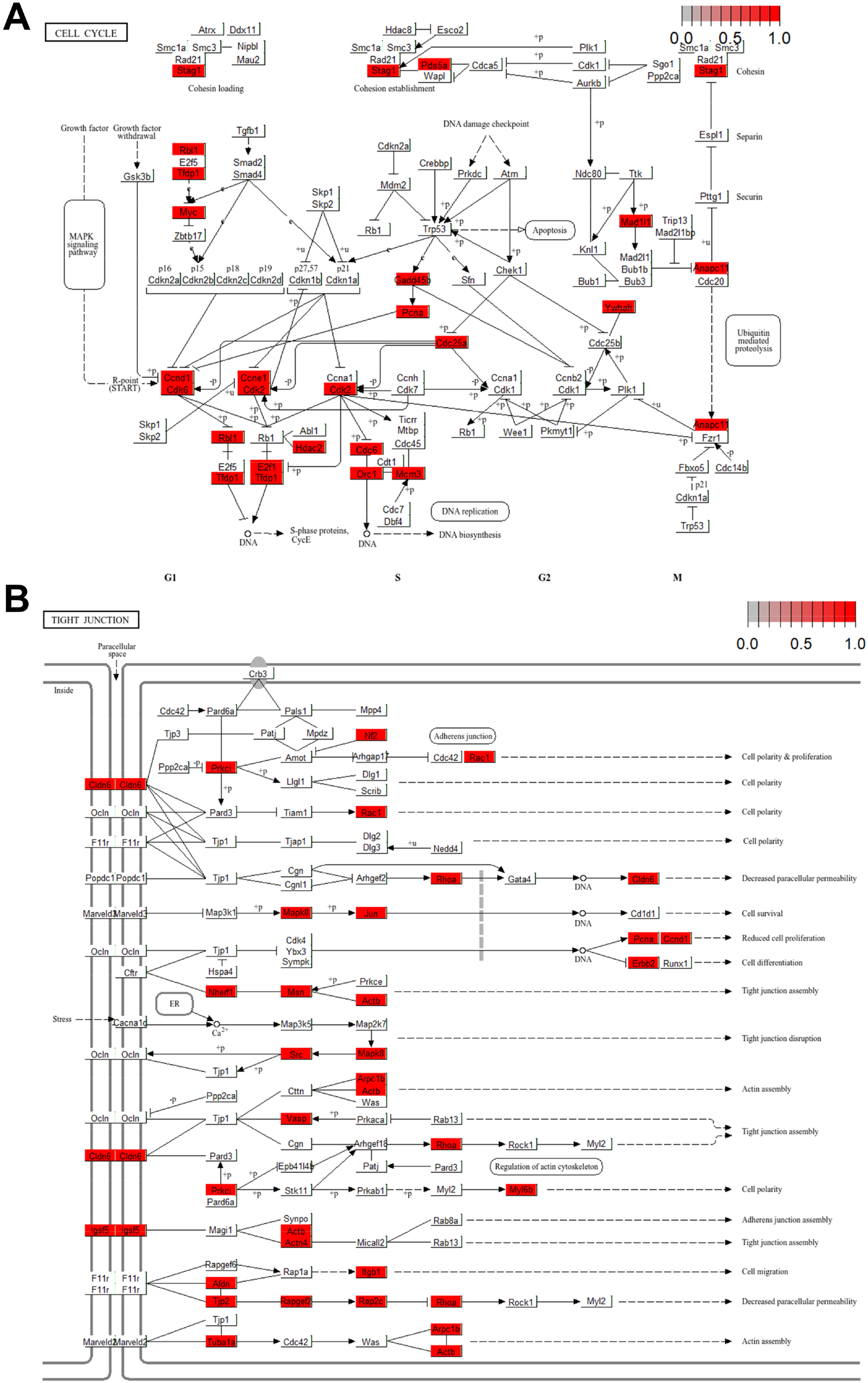
MC38 colon carcinoma cells (MC38^par^, n = 5) or *Ccne2* deleted MC38 cells (MC38*^Ccne2^*^−/−^, n = 5) were cultured for 24 h to allow cells to adhere, before replacing the medium with DMEM containing DMSO (1:1,000) for 20 h followed by RNA isolation, bulk RNA-sequencing and differential gene expression analysis. Over-representation analysis of upregulated genes was performed on KEGG pathways. Schematic representations of (**A**) the “Cell cycle”- and (**B**) the “Tight junction”-KEGG pathway were generated using Pathview with significant up-regulated genes highlighted in red boxes.

**Supplementary Figure 3:**
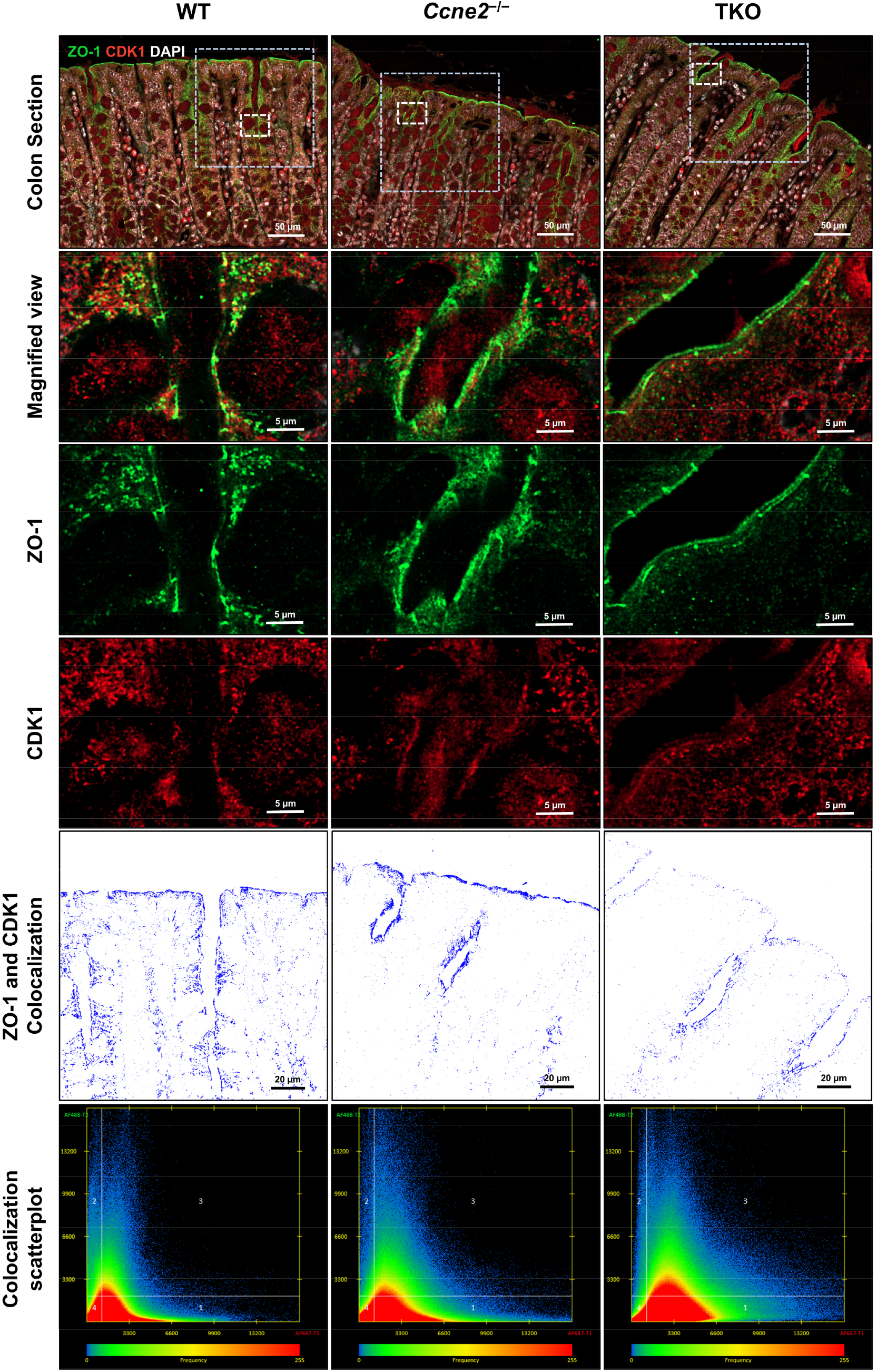
Confocal immunofluorescence microscopy of colon tissue from WT, *Ccne2*^−/−^ and TKO mice, co-stained for ZO-1 (green) and CDK1 (red) with DAPI stained nuclei (white). White dashed boxes in the upper panel correspond to enlarged views shown below; blue dashed boxes correspond to the areas selected for colocalization analysis of the ZO-1 and CDK1 signals shown in the lower panels. Colocalized pixels are displayed in blue based on the thresholds selected in the colocalization scatter plots shown below.

**Supplementary Figure 4:**
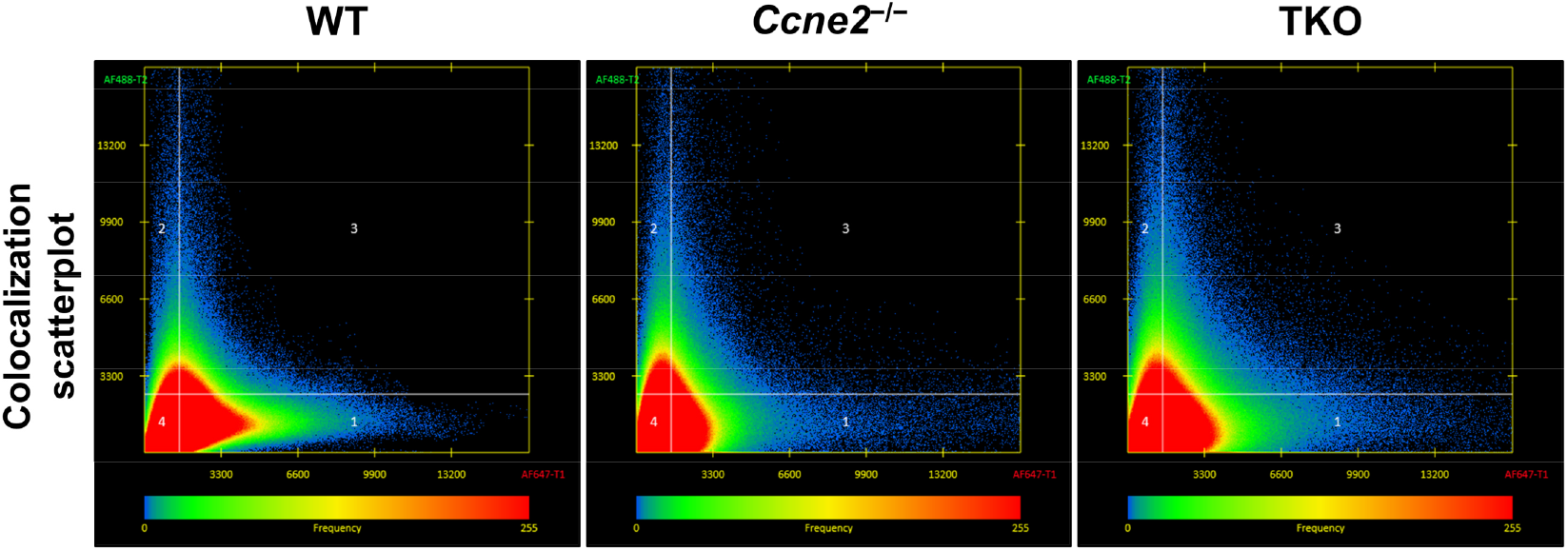
Colocalization scatter plots showing selection of double positive pixels used for the colocalization analysis in the lower panel of Figure 7D.

**Supplementary Figure 5:**
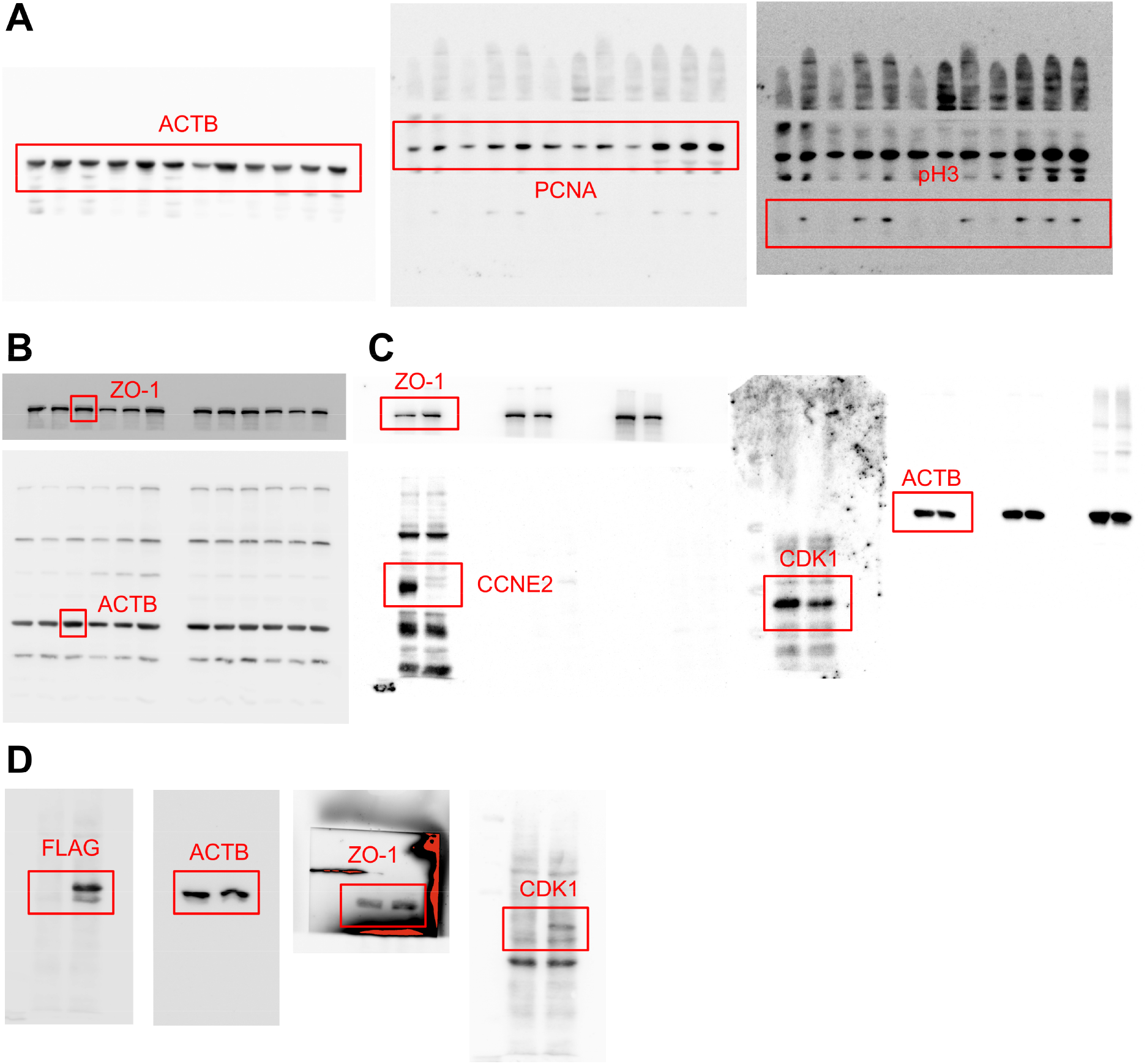
Uncropped Western blots matching to (**A**) Figure 3C, (**B**) Figure 5B, (**C**) Figure 5F and (**D**) Figure 7A. Used lanes are highlighted using red boxes within the raw plots.

**Supplementary Figure 6:**
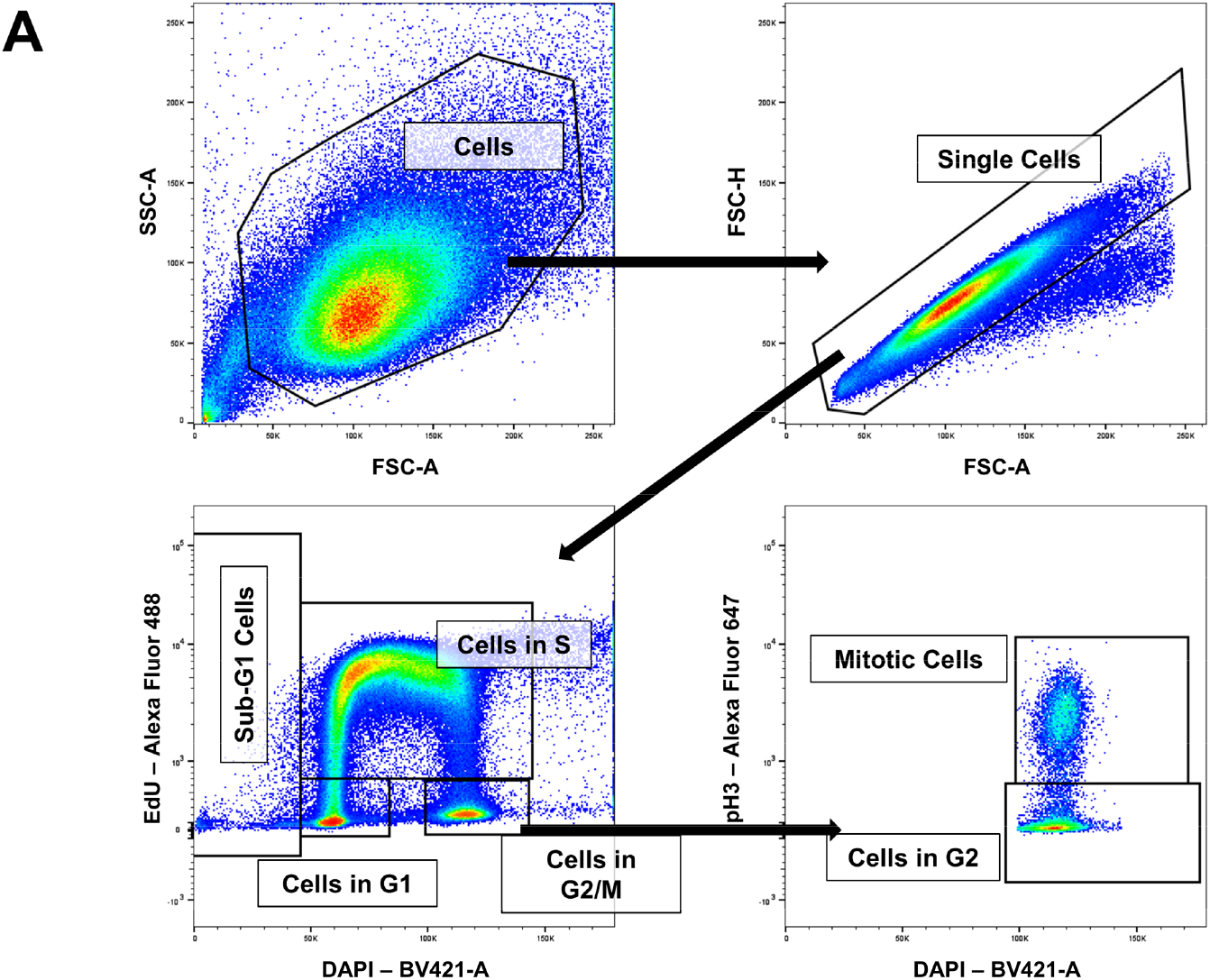
Flow cytometry gating strategy used for cell cycle analysis in Figure 5G-H. Cells were gated based on FSC-A and SSC-A followed by doublet exclusion using FSC-A and FSC-H. Cellular DNA content was measured using DAPI staining and S-Phase activity was measured by EdU incorporation. SubG1 cells were gated based on DNA content lower than 2n which indicates fragmentation of DNA. Cells in G1 phase were based on 2n DNA content with low EdU staining while Cells in S-phase were gated by positive signal for EdU and increasing DNA content. Cells in G2 phase or Mitosis (G2M) were gated based on 4n DNA content and low EdU staining and further subdivided into mitotic cells or cells in G2 phase based on mitosis marker phosphor histone H3 (pH3).

## References

[1] van der Flier LG, Clevers H. Stem cells, self-renewal, and differentiation in the intestinal epithelium. Annu Rev Physiol 2009;71:241–60.

[2] Barker N, van Es JH, Kuipers J, Kujala P, van den Born M, Cozijnsen M, Haegebarth A, Korving J, Begthel H, Peters PJ, Clevers H. Identification of stem cells in small intestine and colon by marker gene Lgr5. Nature 2007;449(7165):1003–7.

[3] Sato T, van Es JH, Snippert HJ, Stange DE, Vries RG, van den Born M, Barker N, Shroyer NF, van de Wetering M, Clevers H. Paneth cells constitute the niche for Lgr5 stem cells in intestinal crypts. Nature 2011;469(7330):415–8.

[4] van de Wetering M, Sancho E, Verweij C, de Lau W, Oving I, Hurlstone A, van der Horn K, Batlle E, Coudreuse D, Haramis AP, Tjon-Pon-Fong M, Moerer P, van den Born M, Soete G, Pals S, Eilers M, Medema R, Clevers H. The beta-catenin/TCF-4 complex imposes a crypt progenitor phenotype on colorectal cancer cells. Cell 2002;111(2):241–50.

[5] Riccio O, van Gijn ME, Bezdek AC, Pellegrinet L, van Es JH, Zimber-Strobl U, Strobl LJ, Honjo T, Clevers H, Radtke F. Loss of intestinal crypt progenitor cells owing to inactivation of both Notch1 and Notch2 is accompanied by derepression of CDK inhibitors p27Kip1 and p57Kip2. EMBO Rep 2008;9(4):377–83.

[6] Bhattacharya S, Ray RM, Johnson LR. Cyclin-dependent kinases regulate apoptosis of intestinal epithelial cells. Apoptosis 2014;19(3):451–66.

[7] Fagundes R, Teixeira LK. Cyclin E/CDK2: DNA Replication, Replication Stress and Genomic Instability. Front Cell Dev Biol 2021;9:774845.

[8] Hu W, Nevzorova YA, Haas U, Moro N, Sicinski P, Geng Y, Barbacid M, Trautwein C, Liedtke C. Concurrent deletion of cyclin E1 and cyclin-dependent kinase 2 in hepatocytes inhibits DNA replication and liver regeneration in mice. Hepatology 2014;59(2):651–60.

[9] Nevzorova YA, Tschaharganeh D, Gassler N, Geng Y, Weiskirchen R, Sicinski P, Trautwein C, Liedtke C. Aberrant cell cycle progression and endoreplication in regenerating livers of mice that lack a single E-type cyclin. Gastroenterology 2009;137(2):691–703, e1-6.

[10] Nevzorova YA, Bangen JM, Hu W, Haas U, Weiskirchen R, Gassler N, Huss S, Tacke F, Sicinski P, Trautwein C, Liedtke C. Cyclin E1 controls proliferation of hepatic stellate cells and is essential for liver fibrogenesis in mice. Hepatology 2012;56(3):1140–9.

[11] Bangen JM, Hammerich L, Sonntag R, Baues M, Haas U, Lambertz D, Longerich T, Lammers T, Tacke F, Trautwein C, Liedtke C. Targeting CCl4 -induced liver fibrosis by RNA interference-mediated inhibition of cyclin E1 in mice. Hepatology 2017;66(4):1242–57.

[12] Penners C, Lin C, Jans AM, Mohs A, Tacke F, Bartneck M, Sonntag R, Liedtke C. The cell cycle regulator Cyclin E1 drives macrophage-associated inflammatory responses in acute immune-mediated hepatitis. bioRxiv 2026:2026.09.30.755646.

[13] Sonntag R, Giebeler N, Nevzorova YA, Bangen JM, Fahrenkamp D, Lambertz D, Haas U, Hu W, Gassler N, Cubero FJ, Muller-Newen G, Abdallah AT, Weiskirchen R, Ticconi F, Costa IG, Barbacid M, Trautwein C, Liedtke C. Cyclin E1 and cyclin-dependent kinase 2 are critical for initiation, but not for progression of hepatocellular carcinoma. Proc Natl Acad Sci U S A 2018;115(37):9282–7.

[14] Turner JR. Intestinal mucosal barrier function in health and disease. Nat Rev Immunol 2009;9(11):799–809.

[15] Tripathi A, Debelius J, Brenner DA, Karin M, Loomba R, Schnabl B, Knight R. The gut-liver axis and the intersection with the microbiome. Nat Rev Gastroenterol Hepatol 2018;15(7):397–411.

[16] Kuo WT, Odenwald MA, Turner JR, Zuo L. Tight junction proteins occludin and ZO-1 as regulators of epithelial proliferation and survival. Ann N Y Acad Sci 2022;1514(1):21–33.

[17] el Marjou F, Janssen KP, Chang BH, Li M, Hindie V, Chan L, Louvard D, Chambon P, Metzger D, Robine S. Tissue-specific and inducible Cre-mediated recombination in the gut epithelium. Genesis 2004;39(3):186–93.

[18] Geng Y, Yu Q, Sicinska E, Das M, Schneider JE, Bhattacharya S, Rideout WM, Bronson RT, Gardner H, Sicinski P. Cyclin E ablation in the mouse. Cell 2003;114(4):431–43.

[19] Odajima J, Wills ZP, Ndassa YM, Terunuma M, Kretschmannova K, Deeb TZ, Geng Y, Gawrzak S, Quadros IM, Newman J, Das M, Jecrois ME, Yu Q, Li N, Bienvenu F, Moss SJ, Greenberg ME, Marto JA, Sicinski P. Cyclin E constrains Cdk5 activity to regulate synaptic plasticity and memory formation. Dev Cell 2011;21(4):655–68.

[20] Ortega S, Prieto I, Odajima J, Martin A, Dubus P, Sotillo R, Barbero JL, Malumbres M, Barbacid M. Cyclin-dependent kinase 2 is essential for meiosis but not for mitotic cell division in mice. Nat Genet 2003;35(1):25–31.

[21] Peterson LW, Artis D. Intestinal epithelial cells: regulators of barrier function and immune homeostasis. Nat Rev Immunol 2014;14(3):141–53.

[22] Jumper J, Evans R, Pritzel A, Green T, Figurnov M, Ronneberger O, Tunyasuvunakool K, Bates R, Zidek A, Potapenko A, Bridgland A, Meyer C, Kohl SAA, Ballard AJ, Cowie A, Romera-Paredes B, Nikolov S, Jain R, Adler J, Back T, Petersen S, Reiman D, Clancy E, Zielinski M, Steinegger M, Pacholska M, Berghammer T, Bodenstein S, Silver D, Vinyals O, Senior AW, Kavukcuoglu K, Kohli P, Hassabis D. Highly accurate protein structure prediction with AlphaFold. Nature 2021;596(7873):583–9.

[23] Tatari MN, De Craene B, Soen B, Taminau J, Vermassen P, Goossens S, Haigh K, Cazzola S, Lambert J, Huylebroeck D, Haigh JJ, Berx G. ZEB2-transgene expression in the epidermis compromises the integrity of the epidermal barrier through the repression of different tight junction proteins. Cell Mol Life Sci 2014;71(18):3599–609.

[24] Slowicka K, Petta I, Blancke G, Hoste E, Dumas E, Sze M, Vikkula H, Radaelli E, Haigh JJ, Jonckheere S, Taminau J, Vandamme N, Wullaert A, Tulchinsky E, Nittner D, Van Vlierberghe P, De Hertogh G, Baldin P, Etlioglu E, Wirapati P, Boon L, Lambrecht BN, Callewaert C, Tejpar S, Goossens S, Berx G, Vereecke L, van Loo G. Zeb2 drives invasive and microbiota-dependent colon carcinoma. Nat Cancer 2020;1(6):620–34.

[25] Terry S, Nie M, Matter K, Balda MS. Rho signaling and tight junction functions. Physiology (Bethesda) 2010;25(1):16–26.

[26] Stauffer S, Zeng Y, Santos M, Zhou J, Chen Y, Dong J. Cyclin-dependent kinase 1-mediated AMPK phosphorylation regulates chromosome alignment and mitotic progression. J Cell Sci 2019;132(20).

[27] Sun X, Yang Q, Rogers CJ, Du M, Zhu MJ. AMPK improves gut epithelial differentiation and barrier function via regulating Cdx2 expression. Cell Death Differ 2017;24(5):819–31.

[28] Santamaria D, Barriere C, Cerqueira A, Hunt S, Tardy C, Newton K, Caceres JF, Dubus P, Malumbres M, Barbacid M. Cdk1 is sufficient to drive the mammalian cell cycle. Nature 2007;448(7155):811–5.

[29] Suski JM, Ratnayeke N, Braun M, Zhang T, Strmiska V, Michowski W, Can G, Simoneau A, Snioch K, Cup M, Sullivan CM, Wu X, Nowacka J, Branigan TB, Pack LR, DeCaprio JA, Geng Y, Zou L, Gygi SP, Walter JC, Meyer T, Sicinski P. CDC7-independent G1/S transition revealed by targeted protein degradation. Nature 2022;605(7909):357–65.

[30] Rutlin M, Rastelli D, Kuo WT, Estep JA, Louis A, Riccomagno MM, Turner JR, Rao M. The Villin1 Gene Promoter Drives Cre Recombinase Expression in Extraintestinal Tissues. Cell Mol Gastroenterol Hepatol 2020;10(4):864–7 e5.

[31] Dantas M, Donker L, Pannekoek WJ, Vliem MJ, van der Beek JA, van Es RM, Vos HR, Klumperman J, Gloerich M. Cyclin-dependent kinase CDK1 targets cell-cell junction components and governs epithelial monolayer integrity throughout the cell cycle. Sci Signal 2026;19(941):eadz1593.

[32] Vandewalle C, Comijn J, De Craene B, Vermassen P, Bruyneel E, Andersen H, Tulchinsky E, Van Roy F, Berx G. SIP1/ZEB2 induces EMT by repressing genes of different epithelial cell-cell junctions. Nucleic Acids Res 2005;33(20):6566–78.

[33] Zhang C, Peng K, Liu Q, Huang Q, Liu T. Adavosertib and beyond: Biomarkers, drug combination and toxicity of WEE1 inhibitors. Crit Rev Oncol Hematol 2024;193:104233.

[34] Guillot A, Kohlhepp MS, Tacke F. Multiplex Immunostaining to Spatially Resolve the Cellular Landscape in Human and Mouse Livers. Methods Mol Biol 2023;2669:245–55.

[35] Meng EC, Goddard TD, Pettersen EF, Couch GS, Pearson ZJ, Morris JH, Ferrin TE. UCSF ChimeraX: Tools for structure building and analysis. Protein Sci 2023;32(11):e4792.

[36] Livak KJ, Schmittgen TD. Analysis of relative gene expression data using real-time quantitative PCR and the 2(-Delta Delta C(T)) Method. Methods 2001;25(4):402–8.

[37] Hubbers A, Hennings J, Lambertz D, Haas U, Trautwein C, Nevzorova YA, Sonntag R, Liedtke C. Pharmacological Inhibition of Cyclin-Dependent Kinases Triggers Anti-Fibrotic Effects in Hepatic Stellate Cells In Vitro. Int J Mol Sci 2020;21(9).

[38] Ewels PA, Peltzer A, Fillinger S, Patel H, Alneberg J, Wilm A, Garcia MU, Di Tommaso P, Nahnsen S. The nf-core framework for community-curated bioinformatics pipelines. Nat Biotechnol 2020;38(3):276–8.

[39] Dobin A, Davis CA, Schlesinger F, Drenkow J, Zaleski C, Jha S, Batut P, Chaisson M, Gingeras TR. STAR: ultrafast universal RNA-seq aligner. Bioinformatics 2013;29(1):15–21.

[40] Patro R, Duggal G, Love MI, Irizarry RA, Kingsford C. Salmon provides fast and bias-aware quantification of transcript expression. Nat Methods 2017;14(4):417–9.

[41] Love MI, Huber W, Anders S. Moderated estimation of fold change and dispersion for RNA-seq data with DESeq2. Genome Biol 2014;15(12):550.

[42] Yu G, Wang LG, Han Y, He QY. clusterProfiler: an R package for comparing biological themes among gene clusters. OMICS 2012;16(5):284–7.

[43] Luo W, Brouwer C. Pathview: an R/Bioconductor package for pathway-based data integration and visualization. Bioinformatics 2013;29(14):1830–1.

[44] Badia IMP, Velez Santiago J, Braunger J, Geiss C, Dimitrov D, Muller-Dott S, Taus P, Dugourd A, Holland CH, Ramirez Flores RO, Saez-Rodriguez J. decoupleR: ensemble of computational methods to infer biological activities from omics data. Bioinform Adv 2022;2(1):vbac016.

[45] Huchzermeier R, Rakateli L, Corcini-Berndt M, Maas SL, Abschlag K, Maassen Pottgens C, Moellmann J, Pirker T, Kremer S, Steitz J, Jin H, Ji B, Dusterhoft S, Ohnmacht C, de Jong RJ, van der Vorst EPC. Deficiency of the aryl hydrocarbon receptor in intestinal epithelial cells, but not hepatocytes, ameliorates atherosclerosis development by sex-dependently targeting the lipid metabolism. Atherosclerosis 2026;420:121857.

[46] Chirumamilla CS, Fazil M, Perez-Novo C, Rangarajan S, de Wijn R, Ramireddy P, Verma NK, Vanden Berghe W. Profiling Activity of Cellular Kinases in Migrating T-Cells. Methods Mol Biol 2019;1930:99–113.

